# Advancing the genetic engineering toolbox by combining *AsCas12a* knock-in mice with ultra-compact screening

**DOI:** 10.1101/2024.05.30.596755

**Authors:** Wei Jin, Yexuan Deng, John E. La Marca, Emily J. Lelliott, Sarah T. Diepstraten, Valentina Snetkova, Kristel M. Dorighi, Luke Hoberecht, Lauren Whelan, Yang Liao, Lin Tai, Geraldine Healey, Wei Shi, Andrew J. Kueh, Benjamin Haley, Jean-Philippe Fortin, Marco J. Herold

**Affiliations:** The Walter and Eliza Hall Institute of Medical Research, Parkville, Melbourne, Australia; Department of Medical Biology, University of Melbourne, Parkville, Melbourne, Australia; Olivia Newton-John Cancer Research Institute, Heidelberg, Melbourne, Australia; School of Cancer Medicine, La Trobe University, Bundoora, Melbourne, Australia; The State Key Laboratory of Pharmaceutical Biotechnology, School of Life Sciences, Nanjing University, Nanjing, China; Department of Molecular Biology, Genentech, Inc., South San Francisco, California, USA; Computational Sciences, Genentech, Inc., South San Francisco, California, USA; current address: Université de Montréal, Centre de recherche de l’Hôpital Maisonneuve-Rosemont

**Author notes:** share first authorship (Wei Jin, Yexuan Deng, John E. La Marca). share senior authorship (Benjamin Haley, Jean-Philippe Fortin, Marco J. Herold).

## Abstract

Cas12a is a gene-editing tool that simplifies multiplexed gene targeting through its RNase activity, enabling maturation of individual crRNAs from a pre-crRNA-encoding RNA. Here, we present a mouse model that constitutively expresses enhanced *Acidaminococcus sp. Cas12a* (*enAsCas12a*) linked to an mCherry fluorescent reporter. We demonstrate efficient single and multiplexed gene-editing in cells from *enAsCas12a^KI^* mice. To test *in vivo* activity, we transduced haematopoietic stem cells from *E*μ*-Myc^T/+^;enAsCas12a^KI/+^*animals with *Trp53*-targeting pre-crRNAs followed by transplantation into irradiated recipient animals. Tumour development was accelerated and TRP53 protein lost. We generated compact, genome-wide Cas12a knockout libraries targeting each gene with four guide RNAs encoded on two (Menuetto) or one (Scherzo) vector. Introducing these libraries into *E*μ*-Myc^T/+^;enAsCas12a^KI/+^*lymphoma cells followed by treatment with an MCL-1 inhibitor (S63845) or TRP53-inducer (nutlin-3a) identified known and novel drug resistance genes. Finally, we demonstrate simultaneous gene knockouts (*Trp53 or combined Bax/Bak*) and activation (*Cd19*) in primary T cells and mouse dermal fibroblasts from crosses of our *enAsCas12a* and CRISPR activation models (*dCas9a-SAM*). Our *enAsCas12a* mouse model and accompanying libraries enhance genome engineering capabilities and complements current CRISPR technologies.

## Introduction

CRISPR-Cas9, the first developed CRISPR-Cas-based gene-editing variant, has found near unparalleled utility in biological and medical research. A particular strength of using CRISPR-Cas9 is the ability to undertake rapid, targeted genetic screens to identify, for example, genes involved in tumourigenesis or clinical drug resistance [1, 2]. Cas12a (Cpf1) is an RNA-guided endonuclease that distinguishes itself from Cas9 by its short CRISPR RNAs (crRNAs), intrinsic RNase activity, and different protospacer adjacent motif (PAM) requirements [3]. RNase activity is necessary for Cas12a to extract mature crRNAs from precursor transcripts (pre-crRNA arrays) by recognition of direct repeat (DR) hairpins upstream (5’) of the crRNA targeting sequence. Concatenating multiple guide RNAs within a single pre-crRNA array is therefore possible, and enables combinatorial targeting of one or multiple genes from a compact, easily-clonable, RNA Pol-III expression cassette [4, 5]. The gene-editing effectiveness of Cas12 has been improved by the engineering of enhanced *Acidaminococcus sp.*-derived *Cas12a* (*enAsCas12a*) [6], but this was previously only applicable in mammalian cells *in vitro*. To extend the applications of Cas12a-mediated gene-editing, we have generated an *enAsCas12a* knock-in transgenic mouse and demonstrated its efficacy when targeting genes individually or simultaneously *in vitro* in primary cells and cell lines, as well as *in vivo* via haematopoietic reconstitution. We have furthermore developed two murine-specific, ultra-compact, pre-crRNA libraries, enabling highly effective genome-scale CRISPR-Cas12a knockout screens in cells derived from this model.

## Results

### Generation and characterisation of the *enAsCas12a* mouse

To introduce Cas12a *in vivo*, we obtained the E174R/S542R/K548R-substitution variant of *Acidaminococcus sp. Cas12a* (*enAsCas12a*; derived form an unclassified *Acidaminococcus* strain (BV3L6)) [6], which has previously been utilised in functional genomic screening [7, 8]. The *enAsCas12a* open reading frame was further modified to contain additional nuclear localisation sequences, which can improve enzyme functionality [9]. The *enAsCas12a* cDNA was then cloned into the Cre-recombinase-inducible expression cassette of a previously described mouse *Rosa26*-targeting construct [10], further modified to include an *IRES-mCherry* marker (instead of *IRES-GFP*), and *enAsCas12a* knock-in (*enAsCas12a^KI^*) mice were then generated by pronuclear microinjection of this construct into *C57BL/6* one-cell stage embryos. We confirmed *enAsCas12a* insertion by long-range PCR (Fig. S1A). Once generated, homozygous *enAsCas12a^KI/KI^*mice were crossed with *CMV-Cre* mice to remove the *loxP*-flanked *neo/stop* cassette and allow constitutive, whole-body *enAsCas12a* expression (Fig. 1A, S1B). This was demonstrated by IRES-mCherry expression in haematopoietic cells/tissues via FACS; almost 100% of peripheral blood cells were mCherry+, and ∼80% of haematopoietic organ cells (thymus, bone marrow, spleen, and lymph nodes) had detectable marker fluorescence (Fig. S1C). To assess enAsCas12a expression-related toxicity, we analysed haematopoietic compartment cell populations, specifically B, T, and myeloid cell types. No changes were observed compared to wildtype controls, suggesting constitutive enAsCas12a expression was well-tolerated (Fig. S2A-D). Furthermore, no health issues were observed in transgenic mice aged up to 250 days (data not shown). Together, these data suggest our *enAsCas12a* mouse model is healthy, and has consistent whole-body expression of the *enAsCas12a* transgene.

**Figure 1.**
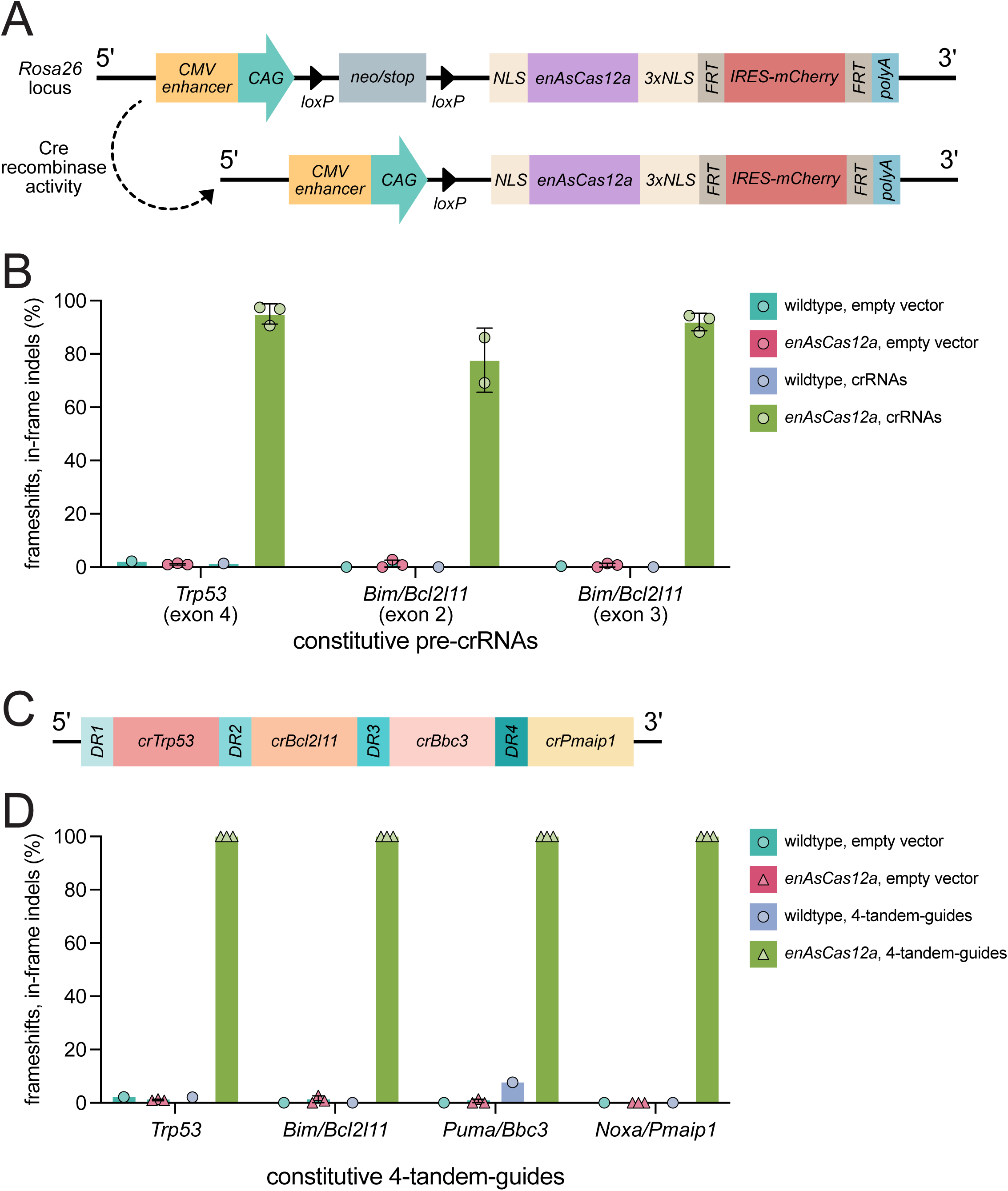
Generation and *in vitro* validation of the *enAsCas12a* knock-in transgenic mouse. (A) Diagram of the *Rosa26*-targeting construct for the genomic insertion of *enAsCas12a*. (B) NGS results showing the efficacy of constitutively expressed crRNAs in MDFs (n=1 WT MDF line, n=3 *enAsCas12a^KI/KI^* MDF lines). (C) Diagram of the 4-tandem-guide construct with pre-crRNAs for parallel targeting of *Trp53*, *Bim/Bcl2l11*, *Puma/Bbc3*, and *Noxa/Pmaip1*. pre-crRNAs are separated by direct repeat sequences (DR1-4). (D) NGS results demonstrate the efficacy of the 4-tandem-guide construct (n=1 WT MDF line, n=3 *enAsCas12a^KI/KI^* MDF lines). In all graphs, the mean is plotted and error bars represent SEM.

### Validation of the *enAsCas12a* gene-editing efficacy *in vitro*

To evaluate enAsCas12a activity, murine dermal fibroblasts (MDFs) from *enAsCas12a^KI^* or WT mice were isolated and transduced with lentiviral vectors for constitutive expression of individual crRNAs targeting *Trp53* (exon 4) or *Bim/Bcl2l11* (exon 2 or 3) (Table S1). Target editing efficiency in transduced *enAsCas12a^KI/KI^*MDFs was ∼100% for each locus, as determined by next generation sequencing (NGS) (Fig. 1B). TRP53 loss was also confirmed via western blotting, with no protein observed even 6 h after treatment with nutlin-3a (MDM2 inhibitor, which leads to indirect TRP53 activation [11]) (Fig S1D). We next adapted our previously described inducible guide RNA expression platform [12], allowing for doxycycline (dox)-mediated temporal control of pre-crRNA activity, for use in combination with our *enAsCas12a*-engineered mouse model. After targeting *Trp53* or *Bim/Bcl2l11 in enAsCas12a^KI/KI^ MDFs*, NGS revealed up to 100% target gene-editing efficiency 24 h post-dox treatment (Fig. S3A, B). At baseline (pre-dox treatment), ∼70-80% of alleles showed signs of editing, representing background leakiness of the inducible promoter.

We then extended our investigations to examine the potential for multiplexed gene-editing via enAsCas12a. A 4-tandem-guide construct was designed for simultaneously targeting *Trp53*, *Bim/Bcl2l11*, *Puma/Bbc3*, and *Noxa/Pmaip1* from a single pre-crRNA expression cassette, with the guides separated by unique DRs on their 5’ end (Fig. 1C). Following lentiviral delivery of this construct, we observed 100% editing efficiency for each targeted gene in *enAsCas12a^KI^* MDFs, and none in WT MDFs (Fig. 1D). NGS also demonstrated 100% gene-editing efficiency for each gene in the dox-treated MDFs using the inducible 4-tandem guide construct. However, as for the inducible single gene pre-crRNAs, leakiness was also observed in pre-dox treated *enAsCas12a^KI/KI^*MDFs (Fig. S3C).

To expand our assessment of enAsCas12a activity to other cell types and biological contexts, homozygous *enAsCas12a^KI/KI^* female mice were crossed with *E*μ*-Myc^T/+^* males, and we generated *E*μ*-Myc^T/+^;enAsCas12a^KI/+^* B lymphoma cell lines from mice that developed MYC-driven lymphoma [13, 14]. We then tested all our constitutive and inducible guides, following lentiviral delivery, in three independent *E*μ*-Myc^T/+^;enAsCas12a^KI/+^*-derived cell lines, and obtained ∼50% gene-editing efficiency for both *Trp53* and *Bim/Bcl2l11* when individually targeted (Fig. S4A, B). Multiplex targeting was more variable, resulting in ∼15% (for the constitutive construct) or ∼35% (for the inducible construct) gene-editing efficiency for *Trp53*, ∼10% (constitutive) or ∼20% (inducible) for *Bim/Bcl2l11*, and less for both *Puma/Bbc3* and *Noxa/Pmaip1* (Fig. S4C, D). In addition, there was less leakage as measured by indel/frameshift mutations pre-dox treatment in *E*μ*-Myc^T/+^;enAsCas12a^KI/+^* cells than in the *enAsCas12a^KI/KI^* MDFs. This suggests that the enAsCas12a system may vary in efficacy based on cell type, or that the presumably higher expression of enAsCas12a in homozygous *enAsCas12a^KI/KI^* MDFs results in more robust activity.

Collectively, these data demonstrate our *enAsCas12a* model is capable of high efficiency single and multiplex gene-editing *in vitro*, using either constitutive or inducible pre-crRNA vectors.

### Validation of the *enAsCas12a* gene-editing efficacy *in vivo*

Having established the efficiency of our enAsCas12a system *in vitro* in both primary and transformed cell lines, our next aim was to evaluate *in vivo* functionality. To do so, we performed haematopoietic reconstitutions, first transducing *E*μ*-Myc^T/+^;enAsCas12a^KI/+^* foetal liver cells (FLCs; obtained at embryonic day 14) with a constitutive *Trp53*-targeting crRNA construct or empty vector control *ex vivo*, then transplanting the transduced FLCs into lethally-irradiated *C57BL/6* recipient mice before monitoring for lymphoma development (Fig. 2A). All mice transplanted with *Trp53*-targeting pre-crRNA-transduced FLCs developed lymphoma by 28 days post-transplantation, (consistent with the latency of reconstituted *E*μ*-Myc^T/+^*;*Trp53^KO^*mice [12, 15]) (Fig. 2B). FACS analyses of the haematopoietic tissues revealed the tumours were primarily immature pro-/pre-B cells (B220^+^IgM^-^IgD^-^) (Fig. 2C). In cell lines derived from the haematopoietic tissues of these mice, successful enAsCas12a-mediated *Trp53* knockout was confirmed by NGS and western blotting analysis (Fig. 2D, E). Collectively, these findings demonstrate that our enAsCas12a system is effective for *in vivo* experimentation.

**Figure 2.**
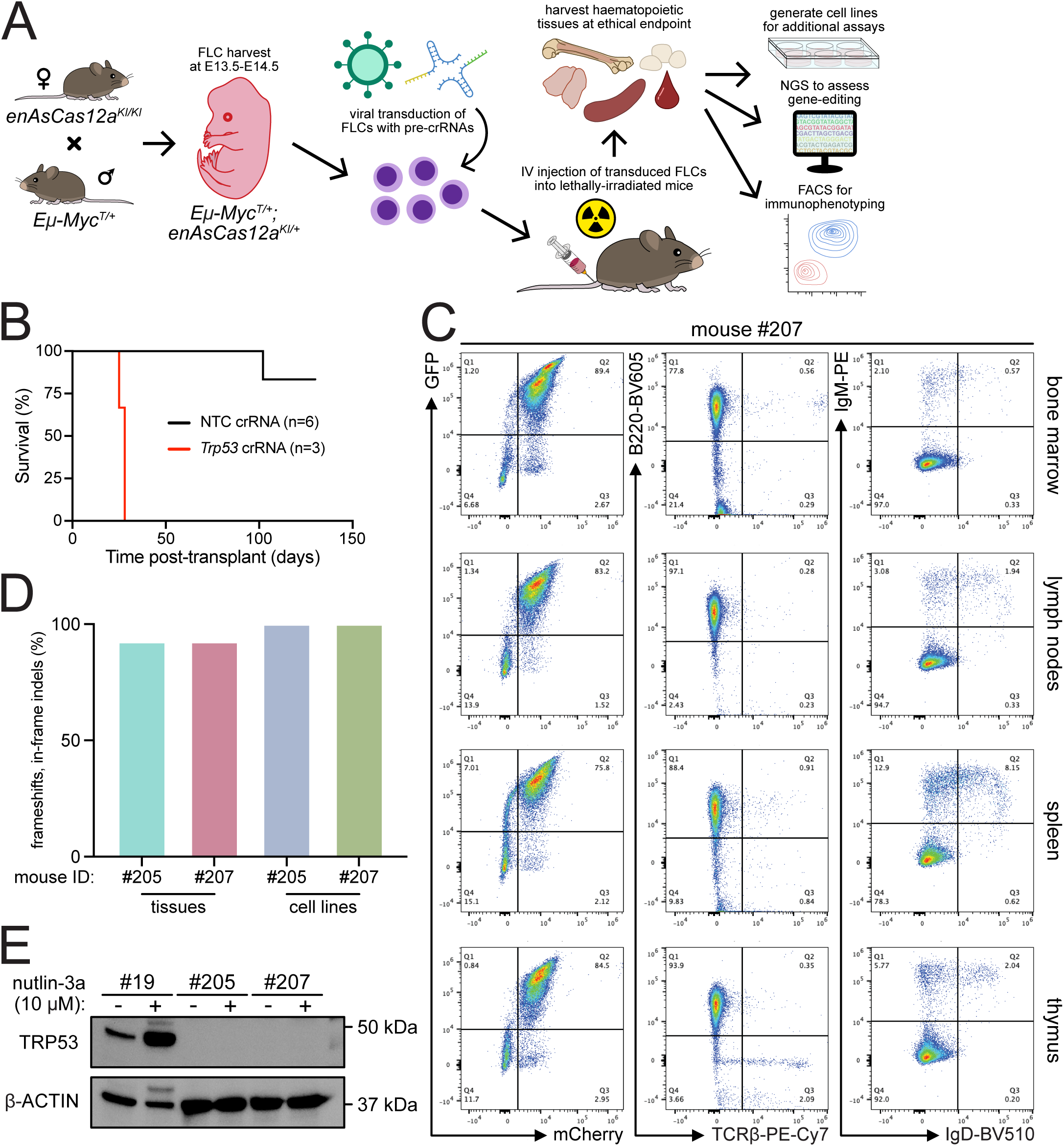
Validation of the enAsCas12a gene-editing efficacy *in vivo*. (A) Schematic of the haematopoietic reconstitution of *E*μ*-Myc^T/+^;enAsCas12a^KI/+^*cells into wildtype animals. (B) Survival curve of mice that underwent haematopoietic reconstitution after transplantation with *E*μ*-Myc^T/+^;enAsCas12a^KI/+^*foetal liver cells, transfected with either a *Trp53*-targeting pre-crRNA (n=3) or an NTC pre-crRNA (n=6). (C) Flow cytometry analysis of haematopoietic organs from a representative mouse (#207, experiment performed for n=3 mice total) reconstituted with the *Trp53*-targeting pre-crRNA *E*μ*-Myc^T/+^;enAsCas12a^KI/+^*foetal liver cells. GFP indicates pre-crRNA presence. (D) NGS results from two representative mice reconstituted with the *Trp53*-targeting pre-crRNA *E*μ*-Myc^T/+^;enAsCas12a^KI/+^*foetal liver cells, demonstrating the efficacy of the knockout. (E) Western blot validating TRP53 loss in two independent *E*μ*-Myc^T/+^;enAsCas12a^KI/+^*;*Trp53^+/-^*cell lines (#205 #207) derived from the splenic tissue of the reconstituted mice. Control cell line #19 was derived from a double-transgenic *E*μ*-Myc^T/+^;enAsCas12a^KI/+^* lymphoma-burdened mouse. TRP53 stabilisation was induced via 24 h treatment with nutlin-3a (in the presence of QVD-O-Ph). β-ACTIN expression was used as a loading control.

### Creation and functional assessment of the Menuetto and Scherzo libraries for whole-genome knockout screening in *enAsCas12a^KI/+^* mouse-derived cells

While several genome-scale crRNA expression libraries compatible with Cas12a have been described for use in human cells (Humagne [7], a “dual” library where there are 4 unique crRNAs per gene across 2 constructs, and Inzolia [8], a “quad” library where 4 unique crRNAs per gene are within a single construct), there are no publicly-disclosed equivalents for screening in murine cells. Therefore, we developed two compact, genome-wide, murine-specific pre-crRNA libraries: Menuetto (dual) and Scherzo (quad). The Menuetto library contains 43,920 constructs with pre-crRNAs targeting 21,743 genes, along with 500 non-targeting controls (NTCs), while the Scherzo library contains 22,839 constructs with pre-crRNAs targeting 21,721 genes plus 500 NTCs (Supplementary File 1).

To demonstrate the utility of the Menuetto and Scherzo libraries, we performed genetic screens using one of our aforementioned *E*μ*-Myc^T/+^;enAsCas12a^KI/+^* cell lines (#20). Six replicate transductions were performed for each library before cells were treated with either DMSO, nutlin-3a (2 μM, retreated 4 times; an MDM2 inhibitor, which leads to indirect TRP53 activation [11]), or S63845 (400 nM, retreated 2 times; an inhibitor of the pro-survival BCL-2 family protein MCL-1 [16]) (Fig. 3A). Drug concentrations close to the IC_50_ values were chosen, as determined via preliminary viability assays (Fig. S5). Once the cells recovered from the multiple treatments, DNA was isolated and NGS was performed to identify enriched crRNAs. Analyses of both the Menuetto (Fig. 3B) and Scherzo (Fig. 3C) library screens revealed similar results: strong enrichment of *Trp53*-targeting pre-crRNAs when comparing nutlin-3a-treated cells to the input samples, and strong enrichment of *Bax*-targeting crRNAs when comparing S63845-treated cells to the input samples. This is in line with expectations, as we have previously identified and validated *Trp53* and *Bax* as resistance factors to nutlin-3a-and S63845-mediated killing, respectively, after conducting whole-genome genetic screens using CRISPR-Cas9 in *E*μ*-Myc* cells [1, 2, 17]. By contrast, the DMSO-treated samples showed no standout guide enrichment for either library (complete analyses of the screens can be found in Supplementary File 2). These data suggest the Menuetto and Scherzo libraries perform well as compact CRISPR-Cas12a whole-genome knockout libraries for use in murine *enAsCas12a* transgenic cells.

**Figure 3.**
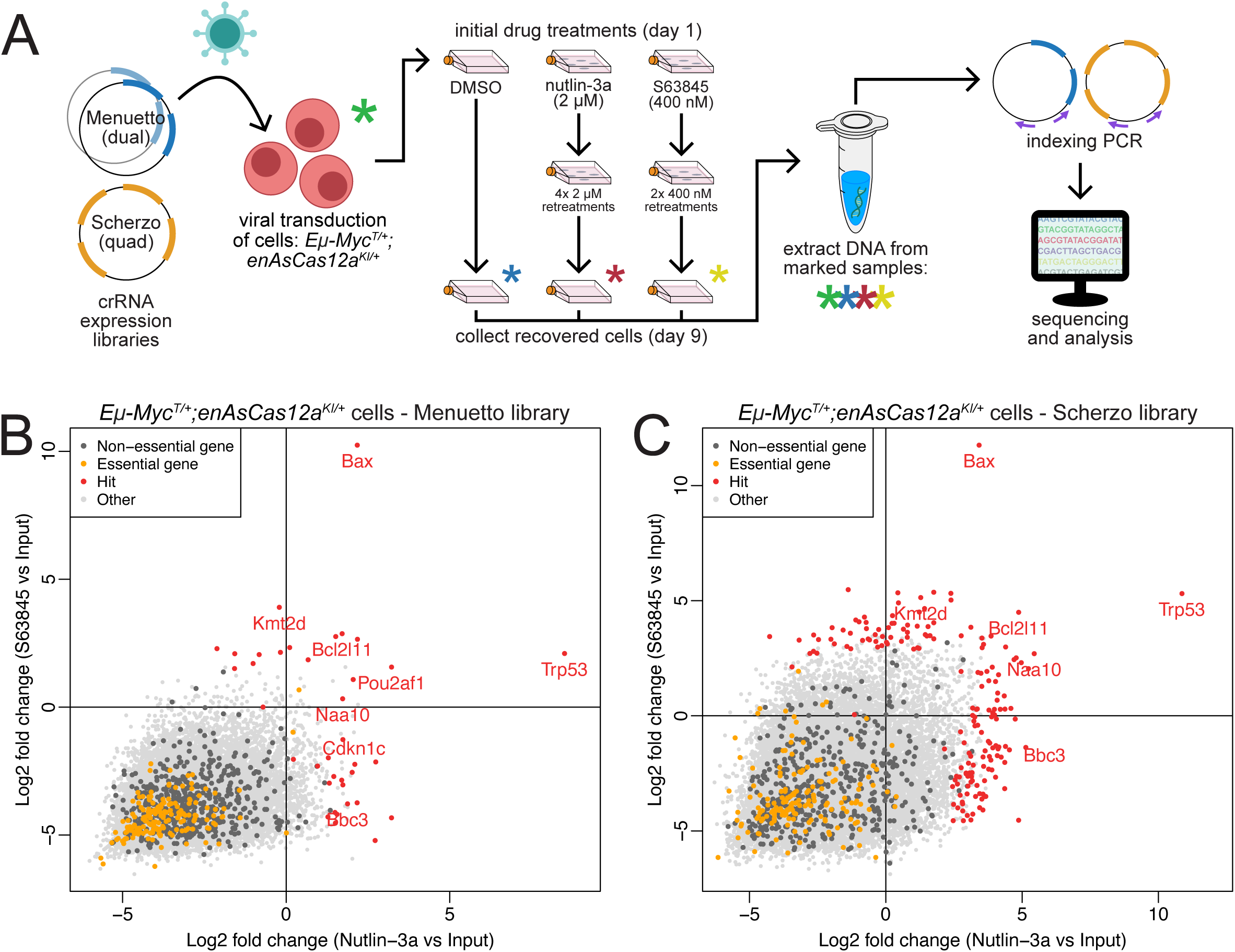
Application of Cas12a ultra-compact, genome-wide, multiplexed murine-specific pre-crRNA libraries. (A) Diagram of the design of the whole genome screens in *E*μ*-Myc^T/+^;enAsCas12a^KI/+^*cells using the Menuetto (dual) and Scherzo (quad) pre-crRNA libraries. Screens of 6 replicate transductions were performed for each library. (B,C) 4-way plots comparing the different arms of the screen samples for the Menuetto (B) and Scherzo (C) libraries. The y-axes compare S63845-treated samples with the input samples, and the x-axes compare the nutlin-3a-treated samples to the input samples. Significantly enriched “hit” genes are indicated in red, essential genes are indicated in orange, non-essential genes are indicated in dark grey, and other genes are indicated in light grey. Complete screen analyses can be found in Supplementary File 2.

### Combinatorial gene modification: *Cas12a*-mediated gene knockout with *dCas9a-SAM*-induced gene expression

Next, we wanted to explore the potential for our *enAsCas12a* mouse to be used for inter-Cas multiplexing applications. In addition to facilitating multiplexed gene knockout through pre-crRNA processing, Cas12a can also be multiplexed with orthogonal Cas molecules, such as a dCas9a used for CRISPR activation (CRISPRa), as their PAM/targeting requirements and guide RNA scaffolds specificities are distinct. To perform such an experiment, we first crossed *enAsCas12a* mice to *OT-I Tg* mice (*OT-I*; in which the T cell receptor is engineered to recognise the immunogen, ovalbumin (OVA) [18]), and the resulting progeny were crossed with our previously described CRISPRa *dCas9a-SAM* mice [19] to generate *enAsCas12a^KI/+^;dCas9a-SAM^KI/+^;OT-I^T/+^* mice. Splenocytes from these mice were stimulated with OVA peptide and transduced with either a *Cd19*-targeting sgRNA (BFP co-expression), a *Trp53*-targeting crRNA (exon 4-targeting; GFP co-expression), or both in combination. Approximately 6% of cells were successfully transduced with both constructs (Fig. 4A, S6A), of which ∼55% expressed CD19, demonstrating successful dCas9a-SAM-mediated *Cd19* activation (Fig. 4B, S6B). We next isolated GFP-negative and GFP-positive cells from the CD19+ population, and used NGS to assess the level of *Trp53* gene-editing. As expected, we observed no genetic perturbations of *Trp53* in CD19+GFP-cells, while *Trp53* was edited with ∼50% efficiency in CD19+GFP+ cells (Fig. 4C, S6C).

**Figure 4.**
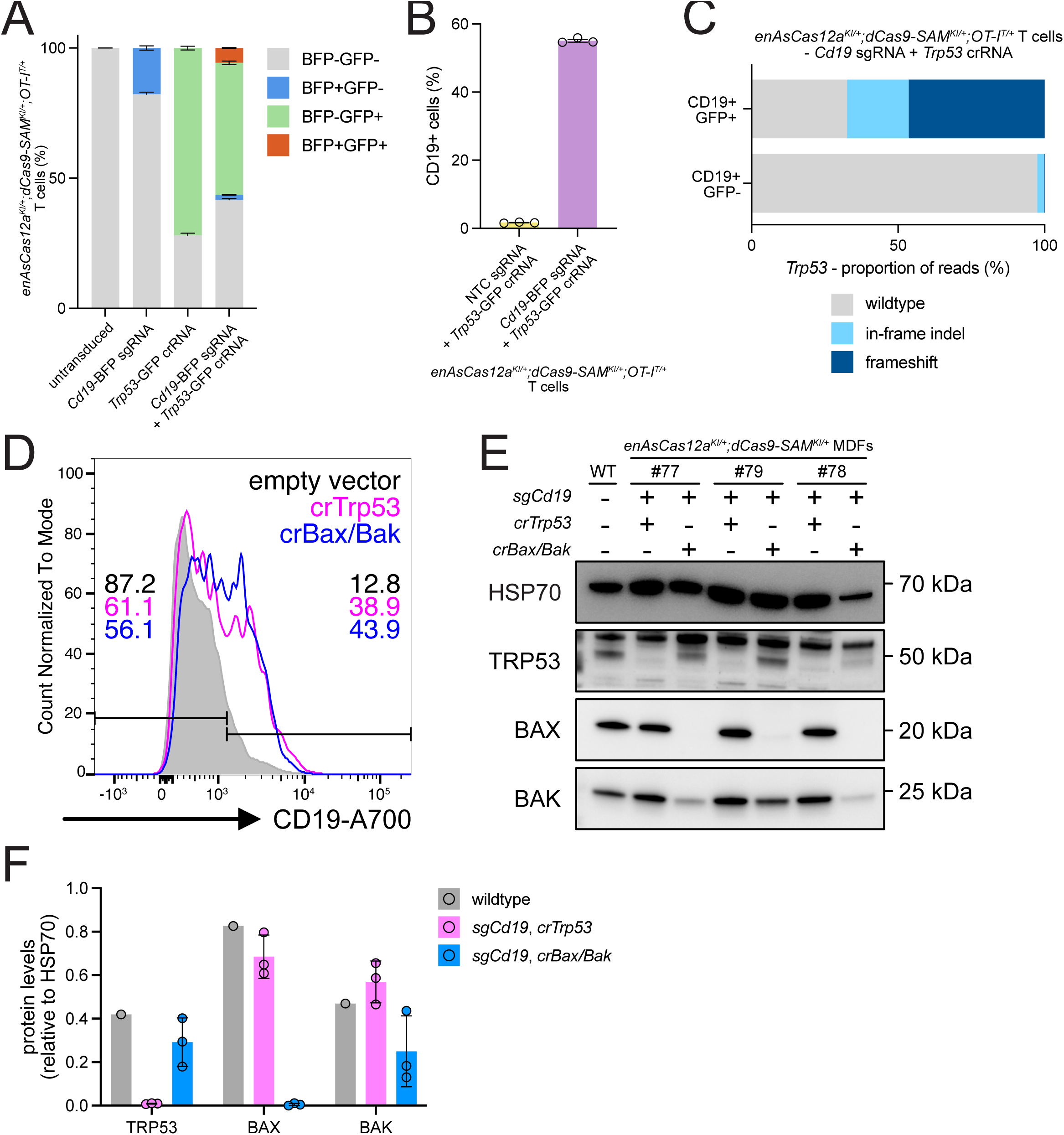
Multiplexing of *enAsCas12a* and *dCas9a-SAM* in *OT-I* T cells. (A) Quantification of FACS analysis of *enAsCas12a^KI/+^;dCas9a-SAM^KI/+^;OT-I^T/+^*T cells transduced with a *Cd19*-targeting sgRNA (BFP-tagged) for dCas9a-SAM-mediated gene expression and a *Trp53*-targeting pre-crRNA (GFP-tagged) for Cas12a-mediated gene-editing. ∼6% of cells from the sample transduced with both constructs were both GFP- and BFP-positive. Results are the averaged data from n=3 transductions. Error bars indicate SD. (B) Quantification of FACS analysis of the T cells transduced with lentiviral vectors expressing the *Trp53*-targeting crRNA and either a NTC sgRNA or the *Cd19*-targeting sgRNA. The *Cd19*-targeting sgRNA was able to induce strong CD19 expression in the double-transduced cells, as determined by intracellular antibody staining. Results represent data from n=3 T cell transductions. In this graph, mean is plotted and error bars represent the SEM. (C) NGS data from the GFP-positive and - negative populations (both CD19-positive) of T cells transduced with both the *Trp53*-targeting pre-crRNA and the *Cd19*-targeting sgRNA. ∼50% of reads indicated a frameshift mutation had been generated in *Trp53*. Sequencing was performed for n=1 transduced T cell line. (D) Representative histogram (normalised to the mode) demonstrating CD19 upregulation in the multiplexed, *sgCd19*-transduced *enAsCas12a^KI/+^;dCas9a^KI/+^*MDFs. CD19 expression for the empty vector transduced MDFs are shown in grey, *crTrp53* in pink, and *crBax/Bak* in blue. (E) Western blots demonstrating TRP53, BAX, and BAK loss in the multiplexed *enAsCas12a^KI/+^;dCas9a^KI/+^* MDFs (n=3). (F) Quantification of the western blots shown in G, demonstrating that while TRP53 and BAX loss were very efficient, BAK loss was incomplete in all samples. In this graph, the means are plotted and the error bars represent SD.

The success of this *enAsCas12a*/*dCas9a-SAM* multiplexing led us to question whether the technique could be performed using a single transduction. To test this, we cloned either the *Trp53*-targeting (exon 4-targeting) crRNA or an array containing both *Bax*- and *Bak*-targeting crRNAs into the same vector as the *Cd19*-targeting sgRNA. Then, we transduced these vectors into *enAsCas12a^KI/+^;dCas9a^KI/+^* MDFs (or the empty vector as a control), and examined gene activation efficacy by flow cytometry 8 days post-transduction. FACS analysis first demonstrated that the majority of our transduced *enAsCas12a^KI/+^;dCas9a^KI/+^* MDFs were triple-fluorescent (mCherry co-expression from *enAsCas12a*, GFP co-expression from *dCas9a-SAM*, and BFP co-expression from the crRNA/sgRNA vector), a useful property for multiplexing applications (Fig. S6D). In those triple-fluorescent cells, we were then also able to observe strong CD19 induction relative to those MDFs transduced with the empty vector (Fig. 4D, S6E). We then examined the efficacy of our gene-knockout via western blotting. For each of the *enAsCas12a^KI/+^;dCas9a^KI/+^* MDFs transduced with the *sgCd19/crTrp53* vector, we observed clear loss of TRP53 expression (Fig. 4E, F). Similarly, for *enAsCas12a^KI/+^;dCas9a^KI/+^* MDFs transduced with the *sgCd19/crBax/crBak* vector, we observed clear loss of BAX, and strong (but incomplete) loss of BAK (Fig. 4E, F).

Together, these data demonstrate that multiplexing enAsCas12a with dCas9a-SAM is a viable strategy for simultaneous knockout and activation of distinct target genes.

## Discussion

In this study, we describe the generation of a murine model containing a constitutively-expressed enhanced *Acidaminococcus sp.*-derived *Cas12a* construct (*enAsCas12a*). We were able to validate that constitutive enAsCas12a expression is well-tolerated, particularly with respect to the haematopoietic system. Furthermore, enAsCas12a expression is present in all surveyed tissues through functional assays and detection of a linked mCherry reporter gene. Efficacy of *enAsCas12a* was determined both *in vitro* – using MDF and *E*μ*-Myc* pre-B/B cell lymphoma cell lines with constitutively and inducibly expressed pre-crRNAs – and *in vivo* – using mice haematopoietically reconstituted with *E*μ*-Myc^T/+^;enAsCas12a^KI/+^*foetal liver cells transduced with constitutively expressed pre-crRNAs. Differences in the efficacy of gene perturbation activity across assayed contexts could likely be explained by *enAsCas12a* dosage, either due to the genetic backgrounds (homozygous *vs* heterozygous), cell type-dependent differences with the CAG promoter driving *enAsCas12a*, or some level of selection favouring the edited cells (e.g. *Trp53* loss inhibiting cell death). We further designed and synthesised two whole-genome, multiplexed, mouse-specific Cas12a crRNA expression libraries: Menuetto (dual) and Scherzo (quad), and demonstrated their utility for *in vitro* screens using *E*μ*-Myc* lymphoma cells. Finally, we crossed our *enAsCas12a* mouse with our previously generated *dCas9a-SAM* mouse [19], and demonstrated the capability for multiplexed experiments with our models, simultaneously utilising both Cas molecules in primary T cells and MDFs to activate CD19 expression and mutate *Trp53* or *Bax/Bak.* Notably, these experiments also demonstrated the viability of our *enAsCas12a* system as being functional in a heterozygous context, meaning it could easily be crossed to disease models to enable broad application amongst the scientific community.

Multiplexing experiments are an emerging area of utility for CRISPR technologies, which exploit the specificity of different Cas molecules to their particular pre-crRNAs/sgRNAs. To utilise this capability to its maximum potential, we designed our *enAsCas12a* mouse with an mCherry marker, making it compatible with our previously generated *dCas9a-SAM* mouse, which has a GFP marker [19]. This design choice should improve quality-of-life for the end users, permitting easy detection and/or enrichment of Cas-/construct-positive cells when needed. Combined with, for example, a BFP-tagged sgRNA/pre-crRNA expression vector, we have demonstrated that one can easily obtain triple-fluorescent (or potentially more) cells with significant utility in FACS and imaging applications.

Due to the RNase capability of Cas12a, multiplexing is also simplified at the level of guides, as multiple crRNAs can be generated from a single RNA molecule and a single promoter. We tested both inducible and constitutively-expressed versions of individual pre-crRNAs and a 4-tandem-guide construct (targeting *Trp53*, *Bim/Bcl2l11*, *Puma/Bbc3*, and *Noxa/Pmaip1* in parallel) *in vitro* using primary *enAsCas12a^KI/KI^* MDFs and *E*μ*-Myc^T/+^;enAsCas12a^KI/+^* lymphoma cells. We saw no loss of gene-editing efficacy in primary MDFs when comparing the constitutive and inducible individual and 4-tandem-guide configurations, but enAsCas12a activity was observed without dox-treatment for both inducible configurations in *enAsCas12a^KI/KI^* MDFs. This leakiness was absent in *E*μ*-Myc^T/+^;enAsCas12a^KI/+^*lymphoma cells transduced with the 4-tandem-guide construct, but there was a marked drop in gene-editing efficiency for those pre-crRNAs distal to their promoter. Previous studies have also reported variable efficiencies in either crRNA cleavage or concomitant gene-editing when using multiplexed Cas12a crRNAs, in various species ranging from bacteria and yeast to human cells [5, 20–22]. As we did not observe a reduction in gene-editing efficiency using the 4-tandem-guide constructs in *enAsCas12a^KI/KI^*MDFs, but lower *Bak* (positioned 3’ of the *Bax*-targeting guide in the *Bax/Bak* crRNA array) gene-editing in the *enAsCas12a^KI/+^;dCas9a^KI/+^* MDFs, this suggests these differences in efficiency are likely dependent on differences in Cas12a expression level.

Multiplexing can also refer to multiple guides targeting a single gene, such as has been the case in the previously generated human-specific Cas12a crRNA expression libraries; Humagne [7] and Inzolia [8]. Humagne uses 2 constructs per gene, each with 2 unique pre-crRNAs (dual), while Inzolia uses only one construct per gene with 4 unique pre-crRNAs (quad). We have taken a similar approach in designing two whole-genome, mouse-specific Cas12a crRNA libraries; Menuetto (dual) and Scherzo (quad). There are advantages and disadvantages to each approach. For example, our dual-style Menuetto library potentially holds more analytical power, as multiple crRNA constructs can be detected per gene, which is necessary for some popular CRISPR screen analysis methods (e.g. MAGeCK [23]) to assign statistical significance. Alternatively, our quad Scherzo library is approximately half the size of the dual Menuetto library, which is advantageous for *in vivo* applications due to the maximisation of gene coverage across a restricted number of guide-expressing cells. Regardless, our screening approach suggests both the Menuetto and Scherzo libraries are highly effective. We obtained a number of expected hits, including *Trp53*, *Bax*, *Puma/Bbc3*, and *Bim/Bcl2l11*, all of which are well-characterised as being necessary for S63845-or nutlin-3a-mediated apoptosis in MYC lymphomas [1, 17, 24, 25]. Other strong hits, such as *Naa10*, are less well characterised, but have been reported elsewhere as necessary for TRP53-mediated apoptosis, as well as having been a hit in a similar CRISPR-Cas9 screen performed by our group [2, 26]. We will make the Menuetto and Scherzo libraries available in the near future to the wider scientific community and look forward to their discoveries.

In conclusion, our resultsdemonstrate the power of Cas12a as an extension of our gene-editing toolbox for engineering sophisticated pre-clinical models and interrogating complex biological pathways.

## Acknowledgments

We thank Le Wang and Ashveen Kaur for general lab assistance. We thank the FACS facilities at both the ONJCRI and WEHI. We thank the BioServices facilities at both the ONJCRI and WEHI. We thank Stephen Wilcox and the WEHI genomics facility. The generation of *enAsCas12a* mice used in this study was supported by Phenomics Australia and the Australian Government through the National Collaborative Research Infrastructure Strategy (NCRIS) program.

## Author contributions

WJ and YD performed all experiments relating to mouse characterisation and enAsCas12a validation. JELM and STD performed all screening experiments. EJL performed all T cell experiments, with assistance from GH. LW performed all live mouse-associated techniques. YL and WS contributed to bioinformatics analyses. VS, KMD, BH, LH, and JPF designed the Menuetto and Scherzo libraries. JPF performed all screening bioinformatics analyses. AJK designed the *enAsCas12a* construct and generated the mouse. LT performed all cloning techniques. MJH designed and supervised the study. WJ, JELM, and YD drafted the manuscript, and JELM revised the manuscript. WJ and JELM prepared the figures. All authors reviewed and approved the manuscript.

## Data availability

Data is available in Supplementary Material.

## Code availability

N/A

## Methods

### Animal strains and husbandry

Care and husbandry of experimental mice was performed according to the guidelines established by The Walter and Eliza Hall Institute Animal Ethics Committee and Austin Health Animal Ethics Committee. Transgenic *E*μ*-Myc*, *enAsCas12a^KI^*, *dCas9a-SAM^KI/+^*, and *enAsCas12a^KI/+^;dCas9a-SAM^KI/+^*mice are maintained on a *C57BL/6-WEHI-Ly5.2* background. *E*μ*-Myc* transgenic mice have been described previously [13, 14]. *C57BL/6-WEHI-Ly5.1* mice used for haematopoietic reconstitutions and *C57BL-6-OT-I Tg* mice [18] (RRID:IMSR, JAX:003831) were obtained from The Walter and Eliza Hall Institute breeding facility (Melbourne, Australia).

### *enAsCas12a* transgene creation and transgenic mouse generation

The *pCAG-enAsCas12a(E174R/S542R/K548R)-NLS(nuc)-3xHA (AAS848)* vector (developed by Keith Joung & Benjamin Kleinstiver [6]) was modified to possess an additional 3x nuclear localization signals (NLS; *3x SV40 NLS*) at the C-terminus of the *enAsCas12a* sequence. This modified *enAsCas12a* sequence (and an upstream spacer sequence) was then cloned into the AscI site of a modified *Rosa26*-targeting CTV vector (obtained from Klaus Rajewsky [10]; modified to remove the *pGK-dTA* sequence and replace the *GFP* with *mCherry*). Inducible *enAsCas12a^KI^*mice were generated by pronuclear injection of this *enAsCas12a Rosa26*-targeting vector (7 ng/μL), *Cas9*-targeting sgRNA (10ng/uL; 5’-CTCCAGTCTTTCTAGAAGAT-3’), and Cas9 protein (50 ng/μL) into *C57BL/6J-WEHI* embryos as previously described [27]. Once generated, heterozygous *enAsCas12a^KI/+^* mice were crossed to *CMV-Cre* deleter mice [28] to remove the *loxP*-flanked *neo/stop* cassette, resulting in the generation of heterozygous mice constitutively expressing *enAsCas12a*, which were bred to produce a homozygous *enAsCas12a^KI/KI^*colony.

### Flow cytometry analyses

To analyse enAsCas12a-IRES-mCherry expression, peripheral blood, bone marrow, thymi, spleens, and lymph nodes were harvested from WT (n=3) and homozygous *enAsCas12a^KI^* (n=3) mice (and processed into single cell suspensions where necessary). Red blood cells in the peripheral blood and spleen were removed by addition of red cell lysis buffer (made in house: ammonium chloride (156 mM), sodium bicarbonate (11.9 mM), EDTA (0.097mM)), before the cells were washed twice with 1x PBS (Gibco #14190144) and centrifugation, before resuspension in FACS buffer (1x PBS, EDTA (5μM) (Sigma-Aldrich #E8008), 5% FBS (Sigma-Aldrich #12007C)).

Cas12a toxicity in the mouse haematopoietic system was determined by first harvesting bone marrow, thymi, spleens, and lymph nodes from WT (n=3) and homozygous *enAsCas12a^KI^* (n=3) mice. Tissues were processed into single cell suspensions, cells were counted using a TC20 Automated Cell Counter (BioRad), then stained with Zombie UV (BioLegend #423107; diluted 1:1000 in FACS buffer) to exclude dead cells. Then cells were washed by centrifugation with 1x PBS, and resuspended in a cocktail of FACS buffer with anti-FCR (made in house; 1:10) and fluorochrome-conjugated antibodies against B220 (RA3-6B2-BV605; 1:200; BioLegend #103244), TCRβ (H57-597-PE-Cy7; 1:400; BioLegend #109222), Mac1 (M1/70-APC-Cy7; 1:400; BD Biosciences #557657), Gr1 (RB68C5-Alexa Fluor 700; 1:400; made in house), IgM (5-1-FITC; 1:400; made in house), IgD (11-26c.2a-BV510; 1:400; BD Biosciences #563110), CD4 (GK1.5-PerCP-Cy5.5; 1:800; BioLegend #100434), CD8 (53.6.7-Alexa Fluor 647; 1:400; made in house). Cells were then incubated on ice for at least 25 m, washed twice by centrifugation with 1x PBS, before resuspension in FACS buffer for analysis. T cells used in multiplexing were prepared similarly as above, and the fluorochrome-conjugated CD19 (ID3-A700; 1:400; made in house) antibody used.

For MDF multiplexing experiments, MDFs were isolated from *enAsCas12a^KI/+^;dCas9a^KI/+^*transgenic mice (#77, #78, #79) and transduced with either a *sgCd19/crBax/crBak* vector, a *sgCd19/crTrp53* vector, or an empty vector (as a control). MDFs were detached from their 6-well plates using cell scrapers (Corning, #3010), placed into FACS buffer, filtered, and washed using 1x PBS and centrifugation. MDFs were then resuspended in FACS buffer with anti-FCR and a fluorochrome-conjugated antibody against CD19 (ID3-A700; 1:400; made in house). After being incubated on ice for 25 m, cells were washed with 1x PBS and resuspended in FACs buffer for analysis. All flow cytometry samples in this study were analysed using either an Aurora (Cytek), FACSymphony A3 (BD Biosciences), or an LSR II (BD Biosciences).

### Small-scale Cas12a pre-crRNA design and construct cloning

The Cas12a individual pre-crRNAs and 4-tandem-guide array targeting *Trp53*, *Bim/Bcl2l11*, *Bbc3/Puma*, and *Pmaip1/Noxa* were designed using Benchling (sequences given in Table S1) and ordered from Integrated DNA Technologies (IDT). Individual pre-crRNAs were synthesized with the same DR (5’-TAATTTCTACTCTTGTAGAT-3’) at the 5’ end, which is the sequence recognized by Cas12a, as well as with 4 bp overhangs at the 5’ end for the complementary (5’-TCCC-3’) sequence and reverse complementary (5’-AAAC-3’) sequences, before being cloned into the BsmBI site (vectors had been modified to contain this site) of either the constitutive lentiviral vector FUGW (Addgene #14883) or the doxycycline-inducible vector FgH1tUTG (Addgene #70183). The 4-tandem-guide array was similarly cloned into the BsmBI site of either FUGW or FgH1tUTG.

The *dCas9a-SAM*-compatible *Cd19*-targeting sgRNA vector was generated by first modifying the lenti sgRNA(MS2)_puro optimized backbone (Addgene #73797) to remove the puromycin selection sequence and instead incorporate a BFP marker. The *Cd19* sgRNA sequence (Table S1) was then cloned into the BsmBI site of the modified vector. The sequences encoding the H1 promoter and crRNA targeting *Trp53* or the crRNA array targeting *Bax/Bak* were cloned into the BamHI site of the modified vector.

### Cell culture

Primary murine dermal fibroblasts (MDFs) were isolated from adult mouse tails. The tail skin (dermis and epidermis) was incubated, with agitation, at 4°C for 24 h in 1.5 mL DMEM (Gibco #11995065) with dispase II (2.1 U/mL; Merck #D4693). The dermis was then removed from the epidermis and digested with collagenase IV (0.0408 mg/mL; Merck #C5138) at 37°C for 1 h in 1.5 mL DMEM with 10% foetal bovine serum (FBS; Sigma-Aldrich #12007C). Single cell suspensions were generated by passing digested dermis through a 100 μm sieve/cell strainer (Falcon #3506) into 3 mL of DMEM with 10% FBS before plating into 6 well tissue culture plates (Falcon #352360). MDFs were maintained in culture using DMEM with 10% FBS, passaged using 1x trypsin (Lonza #BE02-007E) to dislodge them from the plates, and were cultured for no longer than 3 weeks.

*E*μ*-Myc^T/+^;enAsCas12a^KI/+^* cell lines #19, #20, and #23 were derived from spleens of double-transgenic *E*μ*-Myc^T/+^;enAsCas12a^KI/+^* mice after they developed lymphoma (euthanised/harvested at the ages of 95, 131, and 105 days old, respectively). *E*μ*-Myc^T/+^;enAsCas12a^KI/+^* cell lines were maintained in cell culture using FMA media [19], and were cultured for no longer than 3 months.

During their transduction process, FLCs were maintained in foetal liver media: α-MEM with GlutaMax (Gibco #32561037), 10% FBS, HEPES (10 mM; Gibco #15630-080), additional GlutaMax (1x; Gibco #35050-061) sodium pyruvate (1 mM; Gibco #11360-070), and β-mercaptoethanol (50 µM; Sigma-Aldrich #M3148), supplemented with mSCF (0.1 µg/mL; Peprotech #250-03), IL-6 (0.01 µg/mL; made in house), TPO (0.05 µg/mL; Peprotech #315-14), and FLT-3 (0.01 µg/mL; made in house) HEK293T cells were used for virus production and were maintained in cell culture using DMEM with 10% FBS.

All cell lines were cultured at 37°C in 10% CO_2_, and were routinely determined to be negative for Mycoplasma infection using a MycoALert detection kit (Lonza #LT07-118).

### Virus production and cell transduction

For transduction of MDFs (50,000 cells per transduction) and *E*μ*-Myc^T/+^;enAsCas12a^KI/+^*(100,000 cells per transduction) cell lines, plasmid DNA (10 μg) was packaged using p-MDL (5 μg), p-RSV-REV (2.5 μg), and p-VSVG (3 μg), using an established calcium phosphate precipitation method [29]. Lentiviral supernatant, with the addition of polybrene (8 µg/mL), was added onto MDFs and centrifuged at 1200xg for 45 m at 32°C. *E*μ*-Myc^T/+^;enAsCas12a^KI/+^*cells being transduced with Menuetto/Scherzo library DNA were centrifuged at 1100xg for 2 h at 32°C.

For transduction of FLCs, plasmid DNA (10 μg) was packaged using p-MDL (5 μg), p-RSV-REV (2.5 μg), and ENV (5 μg). Lentiviral supernatant was first applied to retronectin (32 µg/mL in 1x PBS)-coated tissue culture plates (Thermo Fisher Scientific #150200) by centrifugation, before the FLCs were applied to the plate, as previously described [15].

### Cell sorting and doxycycline-induction

After 2 days of recovery post-transduction, MDFs and *E*μ*-Myc^T/+^;enAsCas12a^KI/+^* cells were sorted using an Aria III (BD Biosciences) to acquire only cells with *enAsCas12a*-IRES-mCherry and pre-crRNA-GFP expression. Cells with inducible constructs were treated with doxycycline (10 μg/mL) for 24 h to induce pre-crRNA expression as per established protocols [12].

### DNA extraction and next generation sequencing

In all instances, genomic DNA (gDNA) was extracted from cells using DNeasy Blood & Tissue kit (QIAGEN #69506). DNA samples were prepared for NGS similarly to previous descriptions [12]. In short, primers that flank one of the pre-crRNA sequences in the 4-tandem-guides construct were designed with 5’ overhangs and ordered from IDT (full primer sequences in Table S2). To amplify each gene region, an initial PCR was performed using gDNA (100 ng) with 1x GoTaq Green Master Mix (Promega #M7123), and 0.5 μM of each primer. Cycling conditions were 18 cycles of 95°C for 2 m, 60°C for 30 s, 70°C for 30 s. To index the samples for sequencing, a second PCR reaction was then performed using the product from the first PCR (1 μL) with 1x GoTaq Green and 0.5 μM of each primer (indexing sequences in Table S3). The cycling conditions were 24 cycles of 95°C for 2 m, 60°C for 30 s, and 70°C for 30 s. From each amplified, indexed PCR product, 5 μL was pooled and purified using 1.0x Ampure Beads (Beckman Coulter #A63880), and the pooled samples were sequenced using a MiSeq (Illumina). Indels were quantified using the CRISPR indel calculator (http://crisprindelcalc.net).

### Haematopoietic reconstitution

Female *enAsCas12a^KI/KI^* mice were crossed with male *E*μ*-Myc^T/+^* mice, and *E*μ*-Myc^T/+^;enAsCas12a^KI/+^*foetal livers from E13.5-14.5 embryos were harvested. Single cell suspensions of FLCs were generated and frozen in freezing medium (90% FBS and 10% DMSO (Sigma-Aldrich #D4540)). *E*μ*-Myc^T/+^;enAsCas12a^KI/+^* FLCs were transduced with non-targeting control (NTC) or cr*Trp53* (targeting exon 4) (Table S1) via spin infection as described above. The next day, these cells were washed using 1x PBS, filtered, then transplanted into lethally-irradiated (two irradiations of 5.5 Gy, ∼3 h apart) 7-8-week-old male *C57BL/6-WEHI-Ly5.1* recipient mice. Tumour-free survival time was defined as the time from FLC transplantation until reconstituted mice reached ethical endpoint post-lymphoma development. At ethical endpoint, peripheral blood was collected and analysed via Advia (Siemens), and tumour tissues were harvested for immunophenotyping by flow cytometry.

### Haematopoietic cell analyses

The bone marrow, thymi, spleens, and lymph nodes from reconstituted mice at ethical endpoint were harvested and processed into single cell suspensions to culture and generate lymphoma cell lines. A small quantity of the single cell suspension was also prepared for FACS analysis, by first being washed twice using 1x PBS, before being stained with ViaDye Red (1:500; Cytek #R7-60008) for 20 m on ice to exclude dead cells. The cells were then washed twice using 1x PBS, and incubated for 25 m on ice in a cocktail of fluorochrome-conjugated antibodies (B220 (RA3-6B2-BV605; 1:200; BioLegend #103244), TCRβ (H57-597-PE-Cy7; 1:400; BioLegend #109222), IgM (5-1-PE; 1:400; made in house), IgD (11-26c.2a-BV510; 1:400; BD Biosciences #563110)) diluted in FACS buffer with anti-FCR, as above. Cells were then washed with and resuspended in FACS buffer, and analysed by flow cytometry using an Aurora (Cytek).

### Western blot analyses

*E*μ*-Myc^T/+^;enAsCas12a^KI/+^;crTrp53* cell lines derived from lymphoma-burdened reconstituted mice (#205 #207) were treated with the pan-Caspase inhibitor QVD-O-Ph (25 μM; MedChemExpress #HY-12305) for 15 min to inhibit cell demolition, before being treated with nutlin-3a (10 μM; MedChemExpress #HY-10029) for 24 h to induce TRP53 activation (control cells were treated with DMSO). MDF samples (except those in the multiplexing experiments) were treated with nutlin-3a (10 μM) for 6 h with no QVD-O-Ph, but otherwise prepared the same way. Cell pellets were collected and resuspended in RIPA lysis buffer (NaCl (300 mM), SDS (0.2%), Triton X-100 (2%), sodium deoxycholate (1%), Tris HCl (100 mM, pH 8.0)) with protease inhibitor (Roche #11836145001), then incubated on ice for 30 m. Protein-containing supernatant was collected after centrifugation at 13000xg for 10 m at 4°C. To quantify protein content, a BCA assay (Thermo Fisher Scientific #23225) was performed, except for the multiplexing experiments where equal cell numbers were used. Then, 25 μg of protein (or all of the sample for the MDF multiplexing experiment across 2 gels) was loaded into a NuPAGE 4∼12% Bis-Tris 1.5 mm gel (Invitrogen #NP0335) and gel electrophoresis performed. Protein was transferred onto a nitrocellulose membrane (Invitrogen #IB23002) according to the manufacturer’s instructions, and the membrane blocked with 5% skim milk powder dissolved in PBS-T (1x PBS with 0.1% Tween-20 (Sigma-Aldrich #P1379)) for ∼1 h at room temperature, and then incubated in primary antibody against P53 (1:2000; Novocastra #NCL-p53-CM5p), b-ACTIN (1:2000; Sigma #A2228), BAX (1:2000; Sigma Aldrich #B9054), BAK (1:2000; Sigma Aldrich #5897), or HSP70 (1:10,000; gift from Dr R Anderson, ONJCRI) (dissolved in PBS-T) at 4°C overnight, with agitation. The next day, the membrane was washed ∼3 times with PBS-T before incubation with HRP-conjugated anti-rabbit (Southern Biotech #4010-05) or anti-mouse (Southern Biotech #1010-05) secondary antibody. The protein bands were visualised by adding Immobilon Forte Western HRP substrate (Millipore #WBLUF0100) on a ChemiDoc XRS+ (BioRad).

### crRNA design for the Menuetto library (dual pre-crRNA library)

For each mouse protein-coding gene found in Ensembl Release 102, we designed 2 pairs of 23mer spacer sequences for the enAsCas12a nuclease based on the enAsCas12a design rules implemented in the crisprVerse [30]. First, we filtered out spacer sequences with at least one of the following characteristics: contains a poly-T stretch, has GC content below 20% or above 80%, or contains a recognition site for the restriction enzymes EcoRI and KpnI used for lentiviral cloning. Spacer sequences were then selected to (1) minimize the number of putative off-targets located in other coding sequences, (2) optimize on-target activity using the enPAM+GB prediction algorithm described in [7], and (3) target the canonical isoform as defined by Ensembl. We required a minimal distance of 25 nucleotides between spacer sequences within a pair to avoid competing of nuclease occupancy, and required at least 50 nucleotides between pairs as well. When possible, spacers for a given gene were chosen across different exons to increase the probability of having a functional knockout. Spacers located in known Pfam domains, as well as in the first 85% of the CDS region, were prioritized. Finally, to avoid the unintended deletion of functional non-coding elements, we constrained each pair of spacers to regions that do not overlap known non-coding elements described in Ensembl Release 102 (miRNAs, tRNAs, lncRNAs, rRNAs, snRNAs, and snoRNAs). We also included 500 pairs of non-targeting controls (NTCs). The final library, named Menuetto, contains a total of 46,242 pairs of spacer sequences (full Menuetto library information can be found in Supplementary File 1). Pre-crRNAs for the Menuetto library (and Scherzo library, see below) were synthesized and cloned (Cellecta) into a vector derived from the pLKO.1 vector (Addgene #10878) with pre-crRNAs under control of the hU6 promoter and BFP under control of the EF-1α promoter.

### crRNA design for the Scherzo library (quad pre-cRNA library)

The design of the 4 spacers per gene for the quad pre-cRNA library, named Scherzo, follows the design of the Menuetto library. The 4 spacers per gene are also constrained to a region that does not overlap known non-coding elements. As a result, there is a small number of genes for which it is not possible to design one quad array that targets all isoforms of a given gene without overlapping non-coding elements. For such cases, we designed an additional quad array to target the remaining isoforms. The final library contains a total of 22,839 quad arrays, including 500 NTC quad arrays (full Scherzo library information can be found in Supplementary File 1).

### Genome-wide Menuetto and Scherzo crRNA library sequencing

Virus production and *E*μ*-Myc^T/+^;enAsCas12a^KI/+^*cell transduction was performed as described above, with 10 μg library DNA used for each of the six independent transductions undertaken (300,000 cells per transduction) for each library. *E*μ*-Myc^T/+^;enAsCas12a^KI/+^* cells used for screening were not sorted/selected. After recovering from transduction, cell replicates were expanded into T75 flasks (Corning #430641), and treated with either DMSO, Nutlin-3a (2 μM, retreated 4 times; an MDM2 inhibitor, which leads to indirect TRP53 activation), or S63845 (400 nM, retreated 2 times; an MCL-1 inhibitor). Starting drug concentrations were chosen to be around the IC_50_ value, as determined by viability assays performed as previously described [2] (Fig. S5). The first drug treatments occurred ∼7 days post-transduction. After recovering from the final treatment, pellets of 4 million cells were collected, washed with 1x PBS, and the DNA extracted as above.

crRNAs were then amplified from 250 ng of DNA using GoTaq Green according to the manufacturer’s instructions, and the following PCR protocol: 3 m at 95°C, [15 s at 95°C, 30 s at 60°C, 30 s at 72°C repeated 35 times], and 7 m at 72°C. The primers used for amplification and indexation of the pre-crRNAs are given in Table S4. PCRs were performed in duplicate for each sample, with each library indexed separately using the same primer combinations. Products were pooled for each library separately, then cleaned up using Ampure XP beads (Beckman Coulter #A63881) and sequenced on a NextSeq 2000 (Illumina) according to the manufacturer’s instructions.

### Statistical analyses for the Menuetto and Scherzo screens

For both screens, reads were mapped to the libraries using MAGeCK v0.5.9 [23] to generate the raw count data. Raw count data were then stored in a standard Bioconductor SummarizedExperiment object [31] and normalized for sequencing depth. We performed a differential abundance analysis for each pair or quad of pre-crRNAs separately using the popular limma-voom approach [32]. Specifically, we fitted a linear model to the log-CPM values for each pre-crRNA array, using voom-derived observation and quality weights. We performed robust empirical Bayes shrinkage to obtain shrunken variance estimates for each pre-crRNA array, and we used moderated F-tests to compute p-values for each of the two-group comparisons of interest.

For the Menuetto screen, we obtained gene-level statistics by aggregating statistics per gene from the each gene-specific pre-crRNA pair. In particular, we used the “fry” gene-set enrichment analysis method implemented in limma, and considered the two pairs targeting a given gene as a “gene set”. This allows the detection of genes that are consistently enriched or depleted for the two pre-crRNA pairs. We applied the Benjamini-Hochberg procedure to obtain an FDR-corrected p-value for each gene. Essential/non-essential genes were determined as previously described [33]. Hits were selected by using an FDR threshold of 20%.

For the Scherzo screen, the quad array p-values were corrected for multiple comparisons using the Benjamini-Hochberg procedure. Hits were selected by using an FDR threshold of 5%.

### Primary mouse T cell culture, transductions, and cell sorting

Virus production was performed as described above, with 60 μg DNA used in the transduction (∼100×10^6^ splenocytes). Non-treated 6-well plates were coated with retronectin (48 μg/well) overnight at 4°C, washed with 1x PBS and blocked with 2% BSA for 30 m, before the addition of filtered viral supernatant. For multiplexing, each virus was generated independently before supernatants were combined (1:1). Virus was bound to retronectin-coated plates by centrifugation (3500 rpm, 2 h, 32°C), and the supernatant removed. *enAsCas12a^KI/+^;dCas9a-SAM^KI/+^;OT-I^T/+^*splenocytes were resuspended at ∼1×10^6^ cells/mL in T cell media (RPMI-1640 (Gibco #11875093) containing 10% FBS (Bovogen #SFBS), sodium pyruvate (Gibco #11360070), non-essential amino acids (Gibco #11140050), HEPES (Gibco #15630130), Glutamax (Gibco #35050061), penicillin/streptomycin (Gibco #15140122)), stimulated with recombinant human IL-2 (100 U/mL; NIH) and SIINFEKL (10 ng/mL; Auspep) and plated onto the virus-coated plates at ∼8×10^6^ cells/well, before being incubated for 72 h at 37°C. Expanded *enAsCas12a^KI/+^;dCas9a-SAM^KI/+^;OT-I^T/+^*T cells were then washed and maintained in T cell media supplemented with 100 U/mL IL-2 at 0.5-1×10^6^ cells/mL. All cell sorting was performed using an Aria II (BD Biosciences).

### Statistical analyses

Statistical analyses (excluding the analysis of the Menuetto/Scherzo screens described above) were performed using Prism (v10.1.1, GraphPad). All statistical analyses performed reflect comparisons between distinct samples, rather than repeated measures. Differences between two groups were determined by Student’s t-test, having been confirmed to conform with normality assumptions prior. Data are presented as means ± standard error of the mean (SEM), and statistical significance between groups is assessed by P-values, denoted by asterisks (*=P<0.05, **=P<0.01, and ns = no significant difference).

## Supplementary Data

**Supplementary Table 1.**
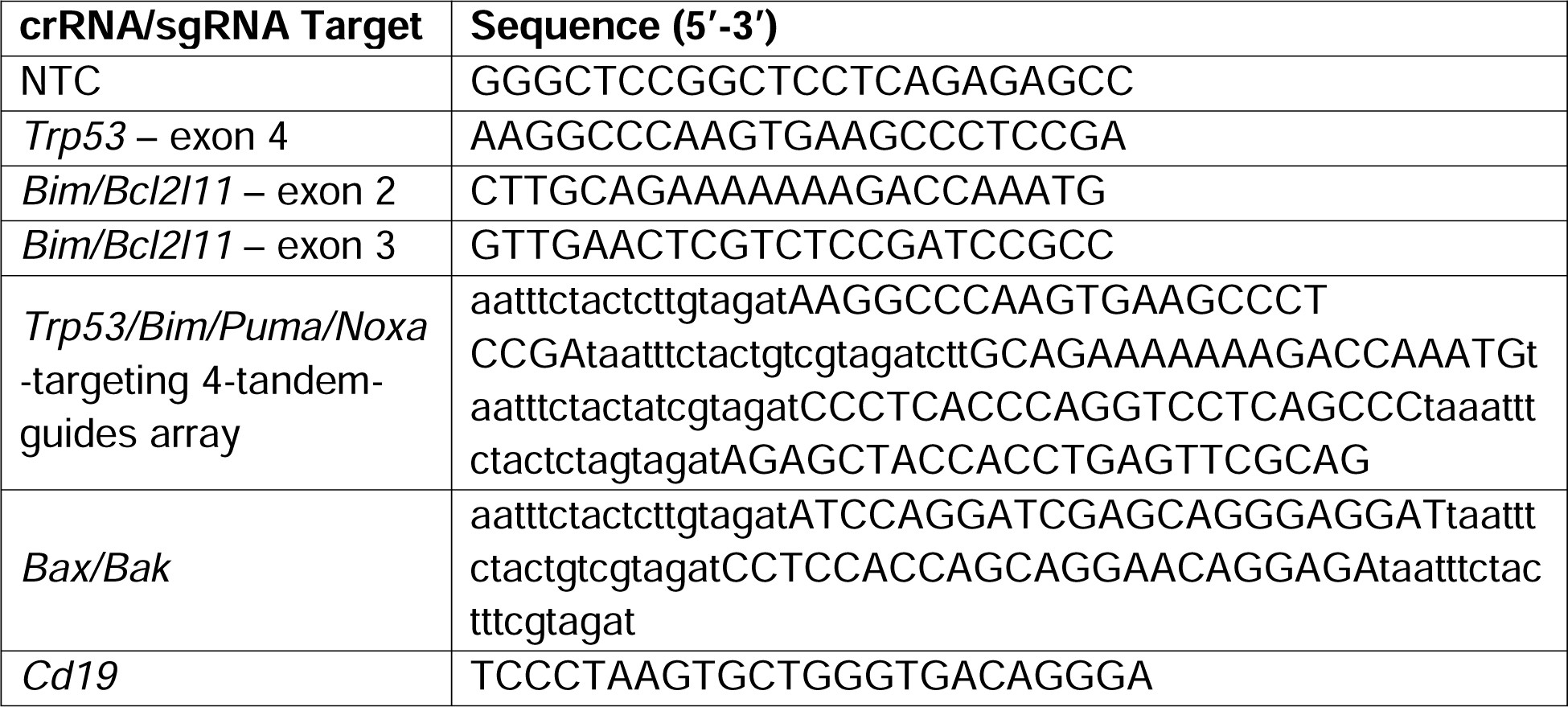
crRNA and sgRNA sequences. Sequences for each crRNA used in this study. The sequence for *Cd19* refers to the *dCas9a-SAM* sgRNA. Note that the lowercase letters in the 4-tandem-guide array sequence refer to the direct repeat sequences.

**Supplementary Table 2.**
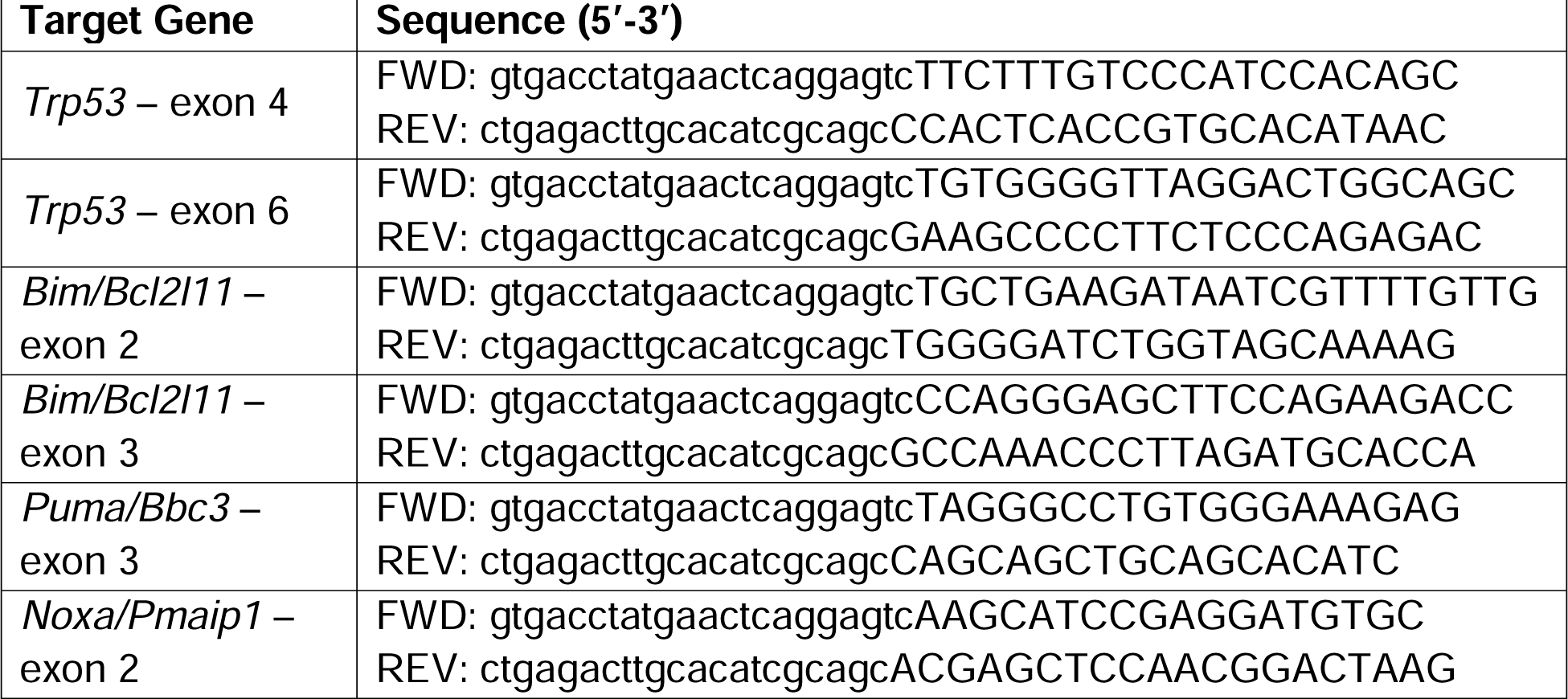
Overhang primers for NGS. Primers used to determine gene knockout efficacy via NGS. Note, the capitalised letters represent the gene targeting sequence, while the lowercase letters represent the overhang sequence.

**Supplementary Table 3.** Indexing primers used in NGS of non-screen samples. Sequences represent the overhang sequences only, and not the entire primer sequence. The remaining primer sequences are complementary to the overhang sequences shown in Table S2.

| Index name (forward) | Index sequence (5'-3') | Index name (reverse) | Index sequence (5'-3') |
| --- | --- | --- | --- |
| Fwd_10 | GAGCTGAA | Rev_01 | TAAGGCGA |
| Fwd_11 | GCGAGTAA | Rev_02 | CGTACTAG |
| Fwd_12 | TGAAGAGA | Rev_03 | AGGCAGA |
| Fwd_13 | TGGTGGTA | Rev_05 | AGGCAGAA |
| Fwd_14 | TTCACGCA | Rev_07 | CTCTCTAC |
| Fwd_15 | AGCACCTC | Rev_08 | CAGAGAGG |
| Fwd_16 | CAAGGAGC | Rev_09 | GCTACGCT |
| Fwd_17 | ATTGGCTC | Rev_10 | CGAGGCTG |
| Fwd_18 | CACCTTAC | Rev_11 | AAGAGGCA |
| Fwd_19 | CTAAGGTC | Rev_12 | GTAGAGGA |
| Fwd_20 | GAACAGGC | Rev_13 | ATGCCTAA |
| Fwd_21 | CCGTGAGA | Rev_14 | ACGCTCGA |
| Fwd_22 | CCTCCTGA | Rev_15 | AGTCACTA |
| Fwd_23 | CGAACTTA | Rev_16 | ATCCTGTA |

**Supplementary Table 4.**
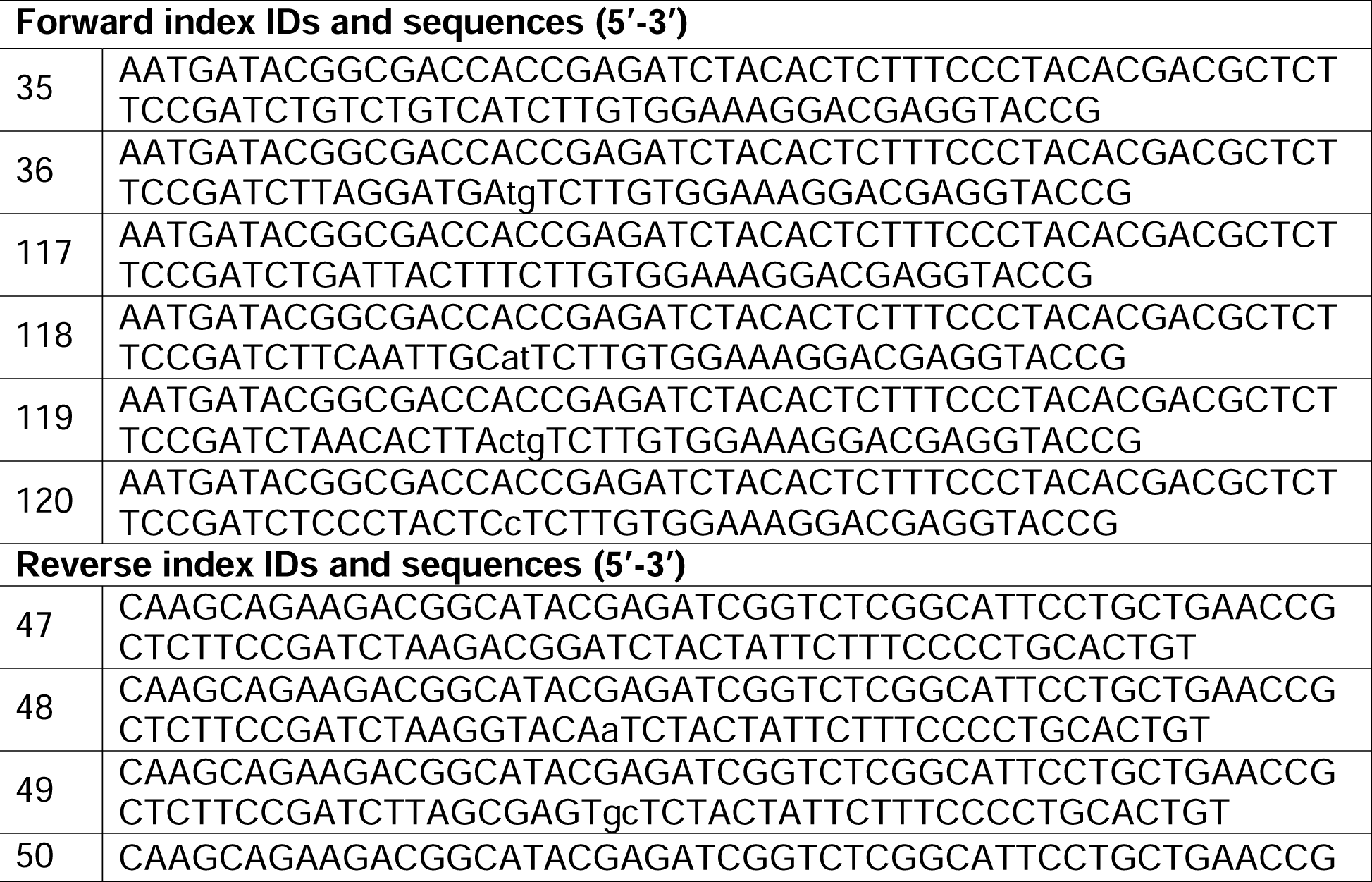
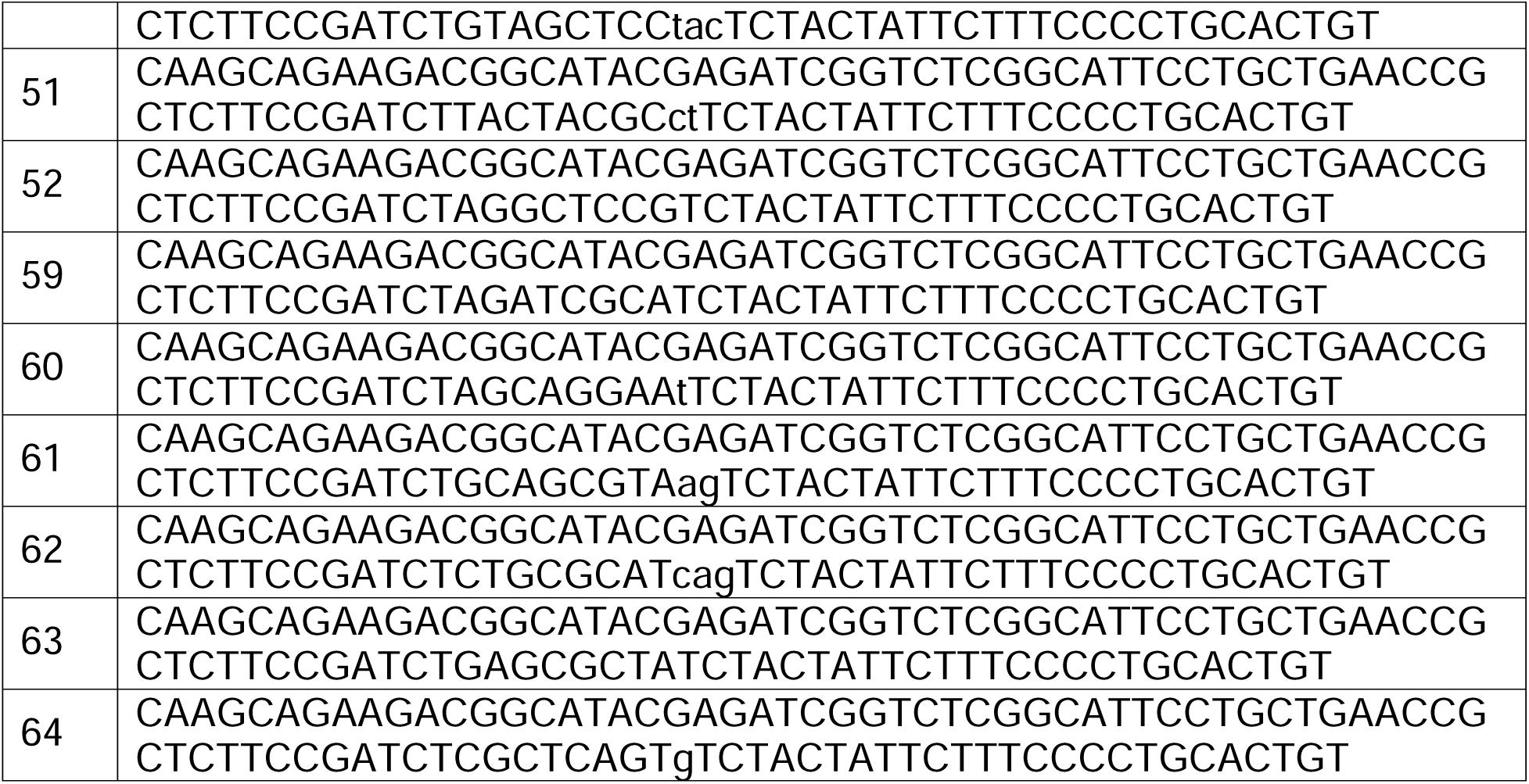
Indexing primers used in NGS of whole-genome screen samples. Entire sequences for the forward and reverse primers used are given.

## Supplementary File Legends

**Supplementary File 1. Cas12a library information.** Complete information for each of the Menuetto and Scherzo Cas12a pre-crRNA libraries, including gene-specific pre-crRNA sequences.

**Supplementary Files 2. Menuetto and Scherzo screen analyses.** DMSO, Nutlin-3a, or S63845 vs input comparisons for each of the whole-genome Cas12a screens conducted. For the Menuetto (dual) library, there are “gRNA” and “gene” level comparisons. For the Scherzo (quad) library, only “gRNA” level comparisons are shown, as this is indistinguishable from the “gene” level due to the design of the library. All comparisons are sorted by FDR.

## Supplementary Figure Legends

**Supplementary Figure 1. Validation of enAsCas12a presence/expression in the transgenic mouse model.** (A) Long range PCR of *enAsCas12a* construct presence in *enAsCas12a^KI/KI^* transgenic mice. The expected band size is 10,126 bp. Samples from 3 *enAsCas12a^KI/KI^* transgenic mice (#40, #41, #42) and 3 control (wildtype (WT)) mice (#76, #81, #83) are shown. (B) The expression of enAsCas12a-linked mCherry in mouse blood was detected by FACS. Data from 1 C57BL/6 WT mouse and 8 enAsCas12a^KI/KI^ transgenic mice (#185, #186, #195∼198, #201, #202) are shown. (C) Haematopoietic organ-specific expression of enAsCas12a-linked mCherry was also detected by FACS. Data from 1 *C57BL/6* WT mouse and 1 *enAsCas12a^KI/KI^* mouse are shown. (D) Western blotting for TRP53, with β-ACTIN expression used as a loading control. MDFs were treated for 6 h with nutlin-3a to induce TRP53 stabilisation. No TRP53 expression was observed in *enAsCas12a^KI/KI^* MDFs with the *Trp53*-targeting pre-crRNA, unlike the WT MDFs or *enAsCas12a^KI/KI^*MDFs with the empty vector.

**Supplementary Figure 2. *enAsCas12a^KI/KI^* mice exhibit no defects in their haematopoietic system.** (A) Representative FACS plots for, and quantification of, T cell distribution between the thymi of *enAsCas12a^KI/KI^* mice (n=3) and wildtype (WT) mice (n=3). (B) Representative FACS plots for, and quantifications of, B cell (pro-B/pre-B = B220^+^IgM^-^; immature B = B220^+^IgM^lo^; mature B = B220^hi^IgM^lo^; transitional B = B220^+^IgM^hi^) and myeloid cell (macrophages = Mac1^+^Gr1^-^; granulocytes = Mac1^+^Gr1^+^) distributions in the bone marrow of *enAsCas12a^KI/KI^* mice (n=3) and WT mice (n=3). (C) Representative FACS plots for, and quantifications of, T cell (TCRβ^+^B220^-^; subsets identified by CD4 and CD8) and B cell (TCRβ^+^B220^+^; subsets identified by IgM and IgD) distribution in the spleens of *enAsCas12a^KI/KI^* mice (n=3) and WT mice (n=3). (D) Representative FACS plots for, and quantifications of, T cell and B cell distributions in lymph nodes of *enAsCas12a^KI/KI^* mice (n=3) and WT mice (n=3), each identified/characterised as above. In each graph, the means are plotted and the error bars represent SD.

**Supplementary Figure 3. Validation of the use of inducible guides in MDFs derived from *enAsCas12a* knock-in transgenic mice.** (A) NGS results showing the efficacy of inducible *Trp53*-targeting crRNAs in WT (n=1) or *enAsCas12a^KI/KI^*(n=3) MDFs. (B) NGS results showing the efficacy of inducible *Bim/Bcl2l11*-targeting crRNAs in WT (n=1) or *enAsCas12a^KI/KI^*(n=3)MDFs. (C) NGS results showing the efficacy of the inducible 4-tandem-guide construct with pre-crRNAs for parallel targeting of *Trp53*, *Bim/Bcl2l11*, *Puma/Bbc3*, and *Noxa/Pmaip1* in WT (n=1) or *enAsCas12a^KI/KI^*(n=3) MDFs. In each graph, the means are plotted and the error bars represent SD.

**Supplementary Figure 4. Validation of enAsCas12a efficacy in *E***μ***-Myc^T/+^;enAsCas12a^KI/+^*cell lines.** (A) NGS results showing the efficacy of constitutively expressed crRNAs targeting *Trp53* or *Bim/Bcl2l11* in *E*μ*-Myc^T/+^*(n=3) or *E*μ*-Myc^T/+^;enAsCas12a^KI/+^* (n=3) cells. (B) NGS results showing the efficacy of inducibly expressed crRNAs targeting *Trp53* or *Bim/Bcl2l11* in *E*μ*-Myc^T/+^*(n=3) or *E*μ*-Myc^T/+^;enAsCas12a^KI/+^* (n=3) cells. (C) NGS results showing the efficacy of the constitutive 4-tandem-guide construct with pre-crRNAs for parallel targeting of *Trp53*, *Bim/Bcl2l11*, *Puma/Bbc3*, and *Noxa/Pmaip1* in *E*μ*-Myc^T/+^* (n=3) or *E*μ*-Myc^T/+^;enAsCas12a^KI/+^*(n=3) cells. (D) NGS results showing the efficacy of the inducible 4-tandem-guide construct with pre-crRNAs for parallel targeting of *Trp53*, *Bim/Bcl2l11*, *Puma/Bbc3*, and *Noxa/Pmaip1* in *E*μ*-Myc^T/+^;enAsCas12a^KI/+^* cells. In each graph, the means are plotted and the error bars represent SD.

**Supplementary Figure 5. Screening preparation.** (A, B) Preliminary viability assays used to determine suitable nutlin-3a (A) and S63845 (B) concentrations used in the Menuetto and Scherzo screens in *E*μ*-Myc^T/+^;enAsCas12a^KI/+^* cells. A dose close to the IC_50_ value was selected for each screen. Viability assays were performed once, with technical replicates performed in duplicate.

**Supplementary Figure 6. Flow cytometry of multiplexed enAsCas12a and dCas9-SAM OT-I T cells.** (A) Representative FACS plots of *enAsCas12a^KI/+^;dCas9a-SAM^KI/+^;OT-I^T/+^*T cells (n=3 total) transduced with different guide RNA constructs. From left to right: untransduced, *CD19*-targeting BFP+ sgRNA for dCas9a-SAM, *Trp53*-targeting GFP+ pre-crRNA for Cas12a, and transduced with both constructs. (B) Representative FACS plots of *enAsCas12a^KI/+^;dCas9a-SAM^KI/+^;OT-I^T/+^*T cells (n=3 total) transduced with a *Trp53* pre-crRNA construct, pre-gated on GFP+BFP+ cells. The cells additionally transduced with a NTC sgRNA show no CD19 expression (left), while those additionally transduced with a *CD19*-targeting sgRNA show strong CD19 expression (right). (C) FACS plots of *enAsCas12a^KI/+^;dCas9a-SAM^KI/+^;OT-I^T/+^* T cells (n=1 total) doubly transduced with *Trp53* pre-crRNA and CD19 sgRNA constructs. CD19-positive cells either GFP-positive or -negative (populations Q2, Q4) were sorted out for NGS to determine *Trp53* editing efficiency. (D) Gating strategy for the multiplexed *enAsCas12a^KI/+^;dCas9a^KI/+^* MDFs (n=3), demonstrating the triple-fluorescent nature of the cells (mCherry co-expression from *enAsCas12a*, GFP co-expression from *dCas9a-SAM*, and BFP co-expression from the crRNA/sgRNA vector). (E) Quantification of the CD19 expression in in the multiplexed *enAsCas12a^KI/+^;dCas9a^KI/+^* MDFs (n=3). A ∼2-3 fold increase in CD19 expression was observed for all samples. In this graph, the means are plotted and the error bars represent SD.

