## Supplementary figures and images for "Advancing the genetic engineering toolbox by combining *AsCas12a* knock-in mice with ultra-compact screening"

### Supplementary Figures

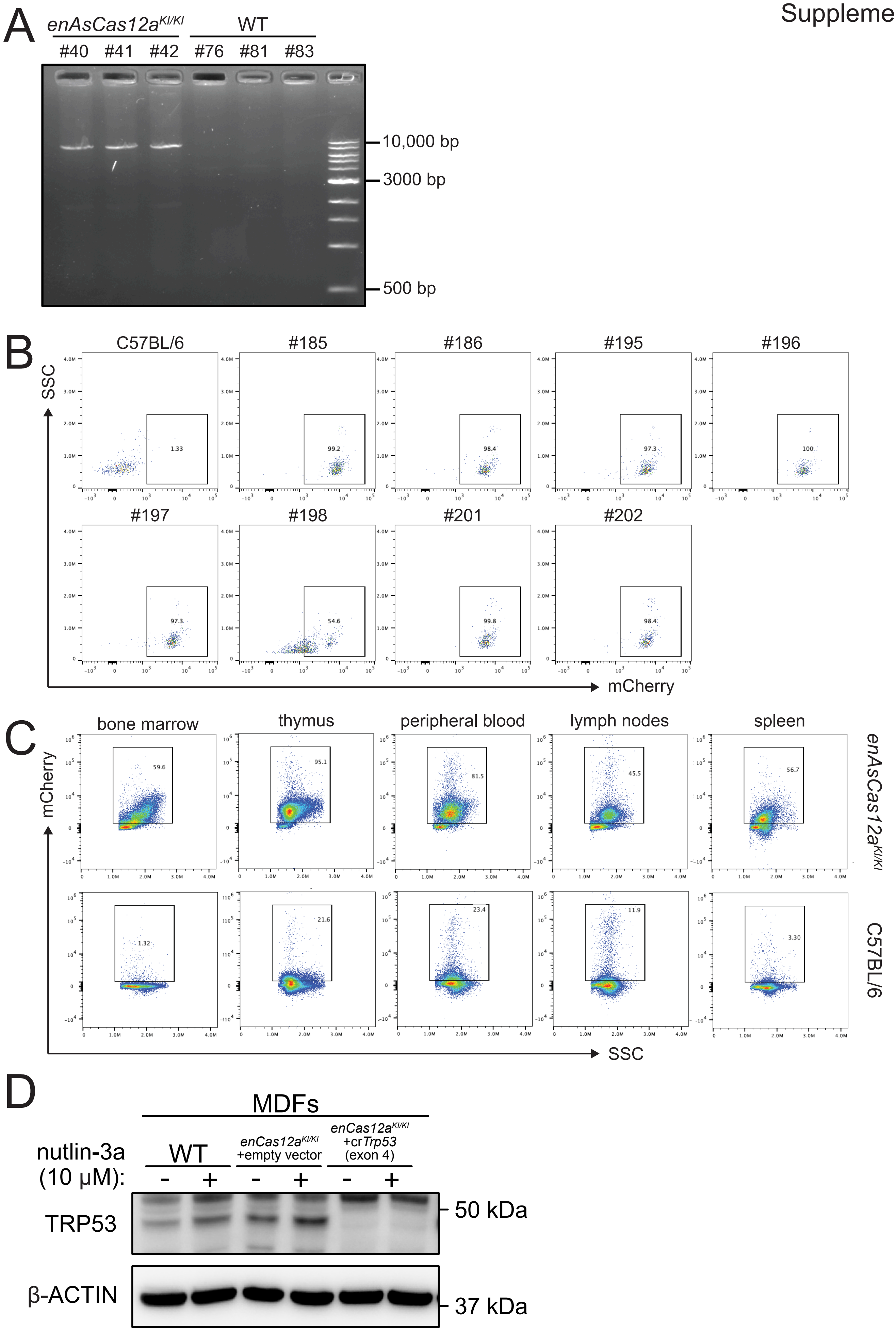

Supplementary Figure 2

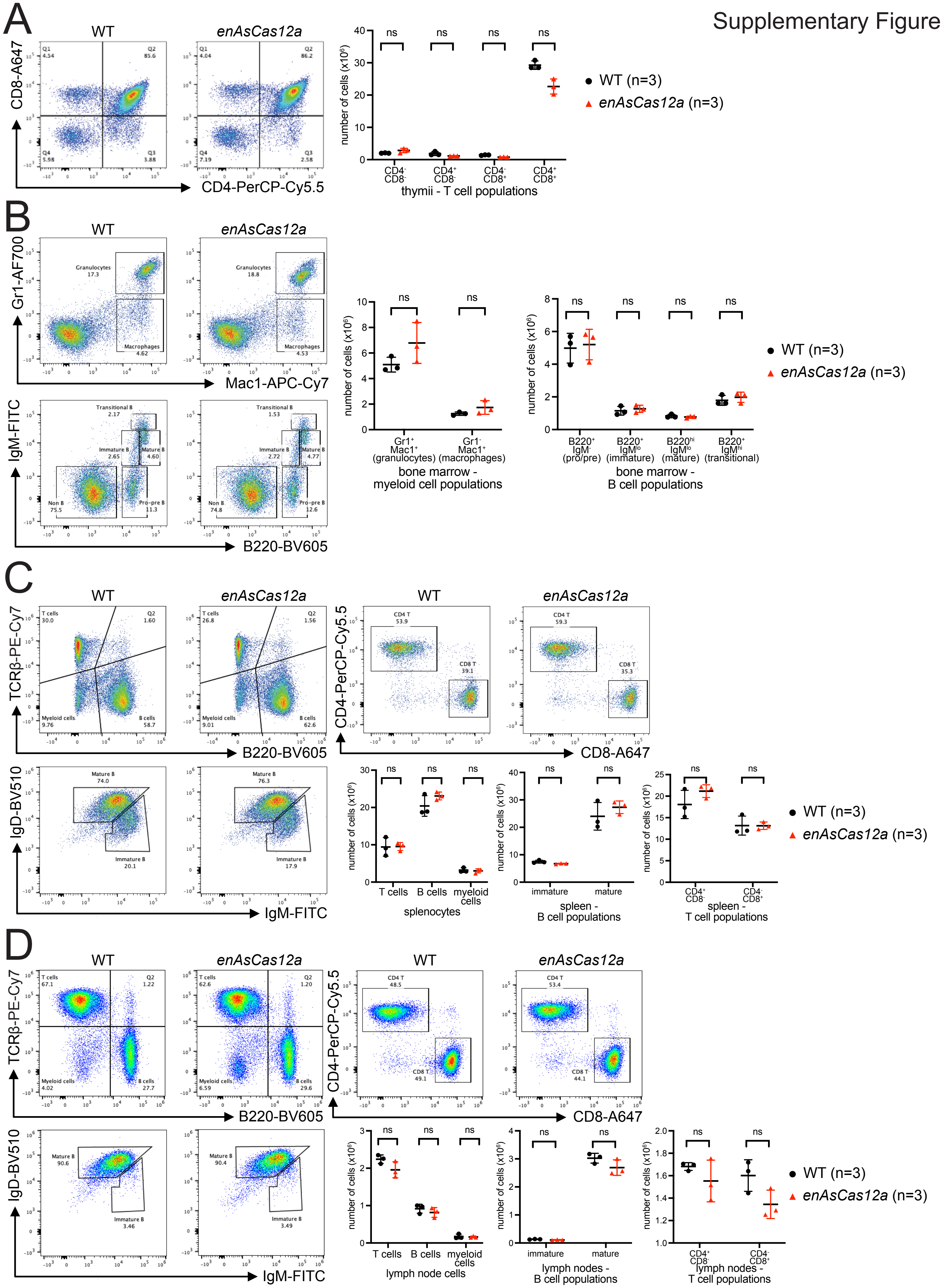

#### Supplementary Figure 3

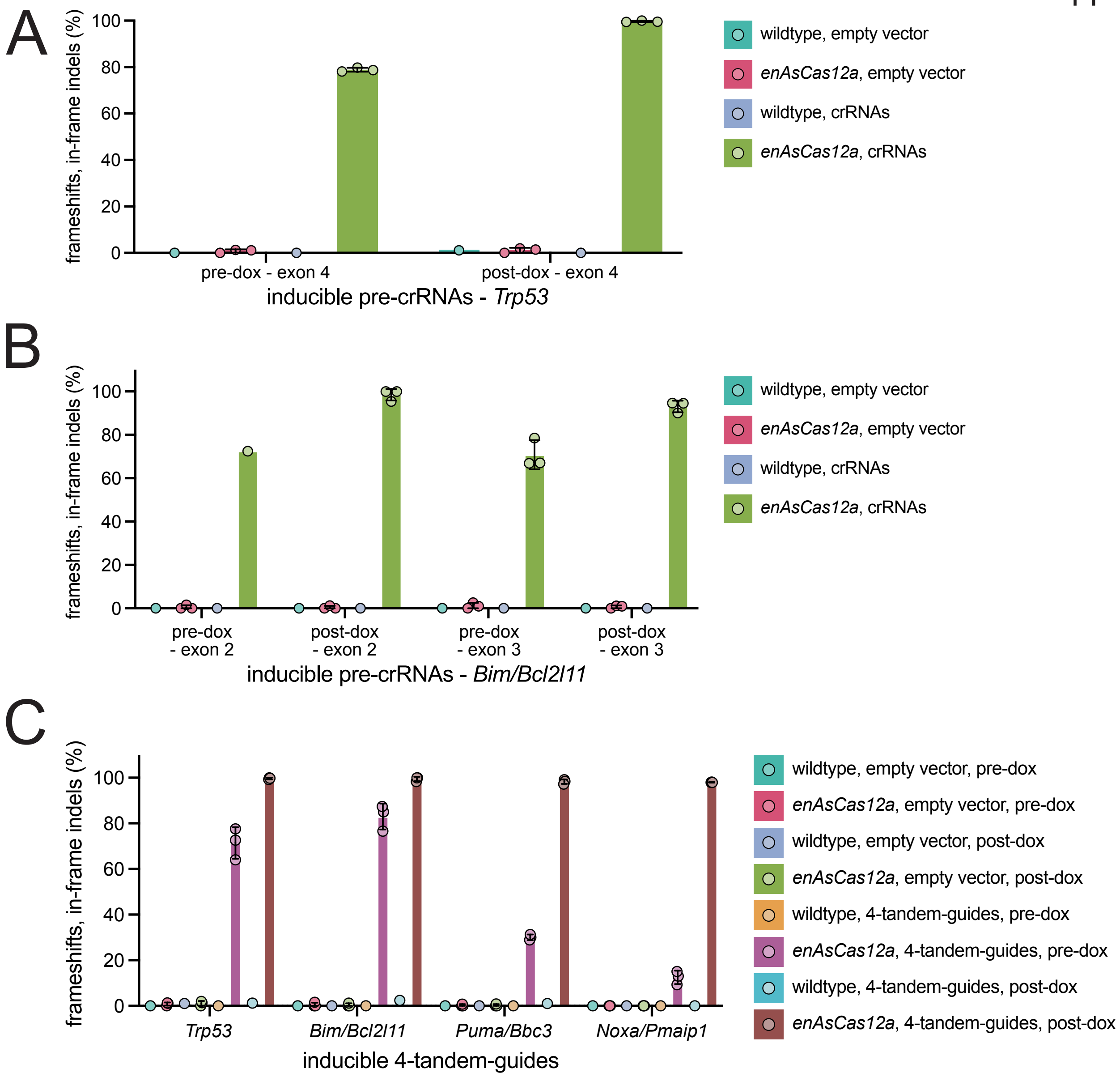

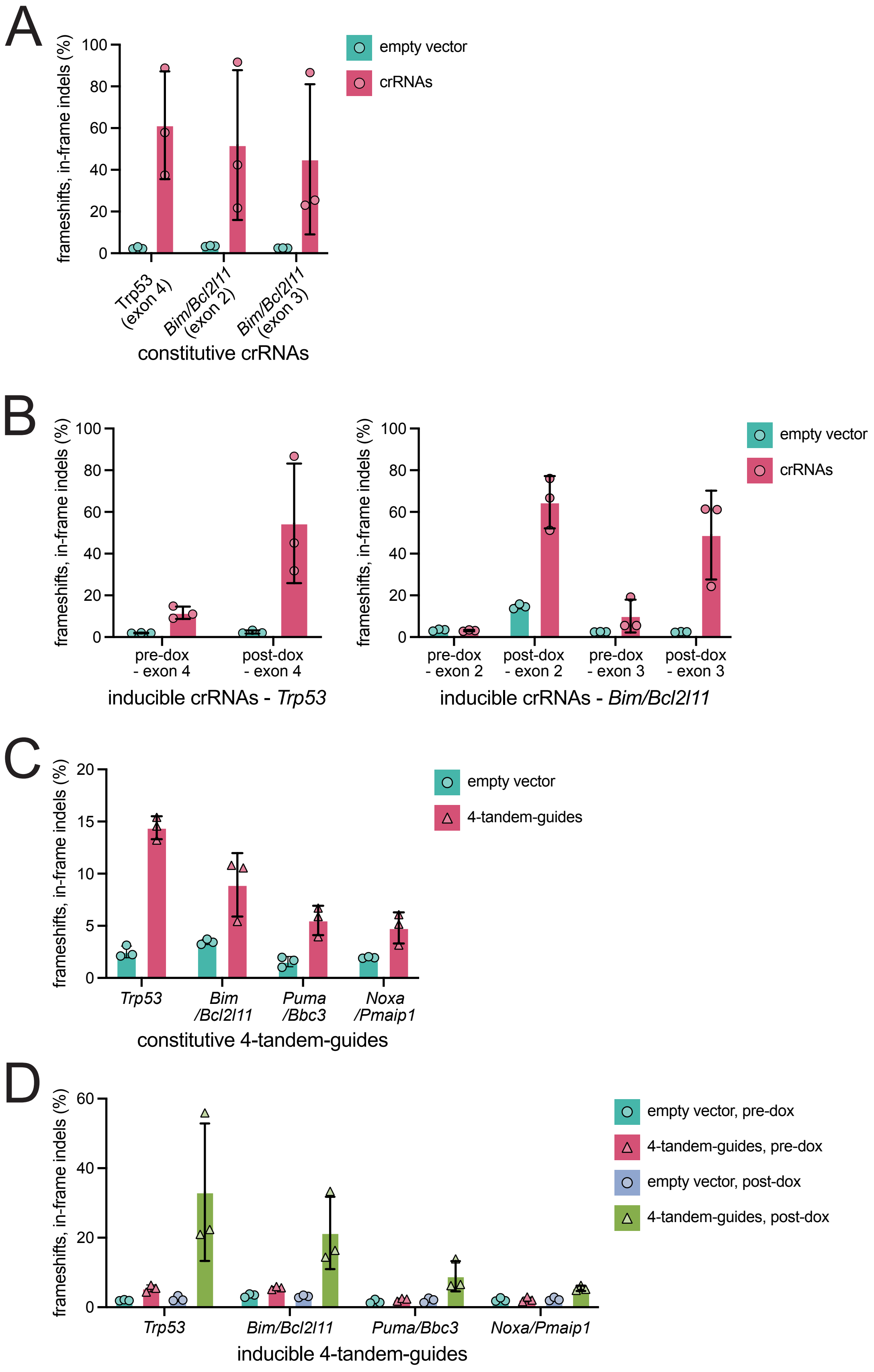

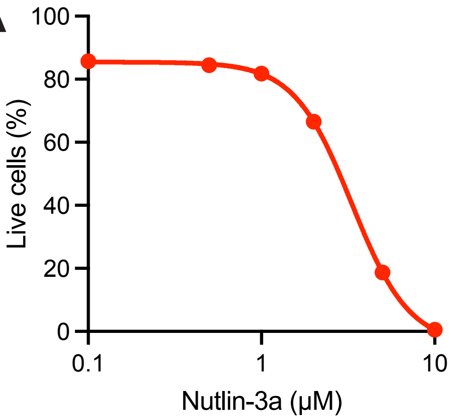

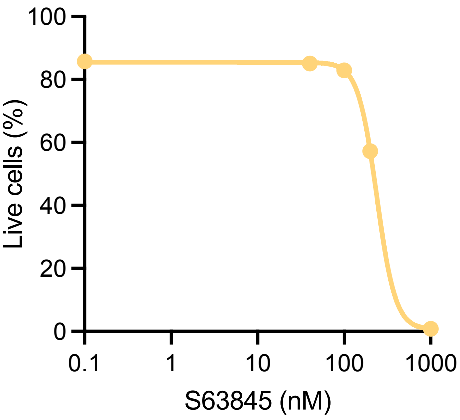

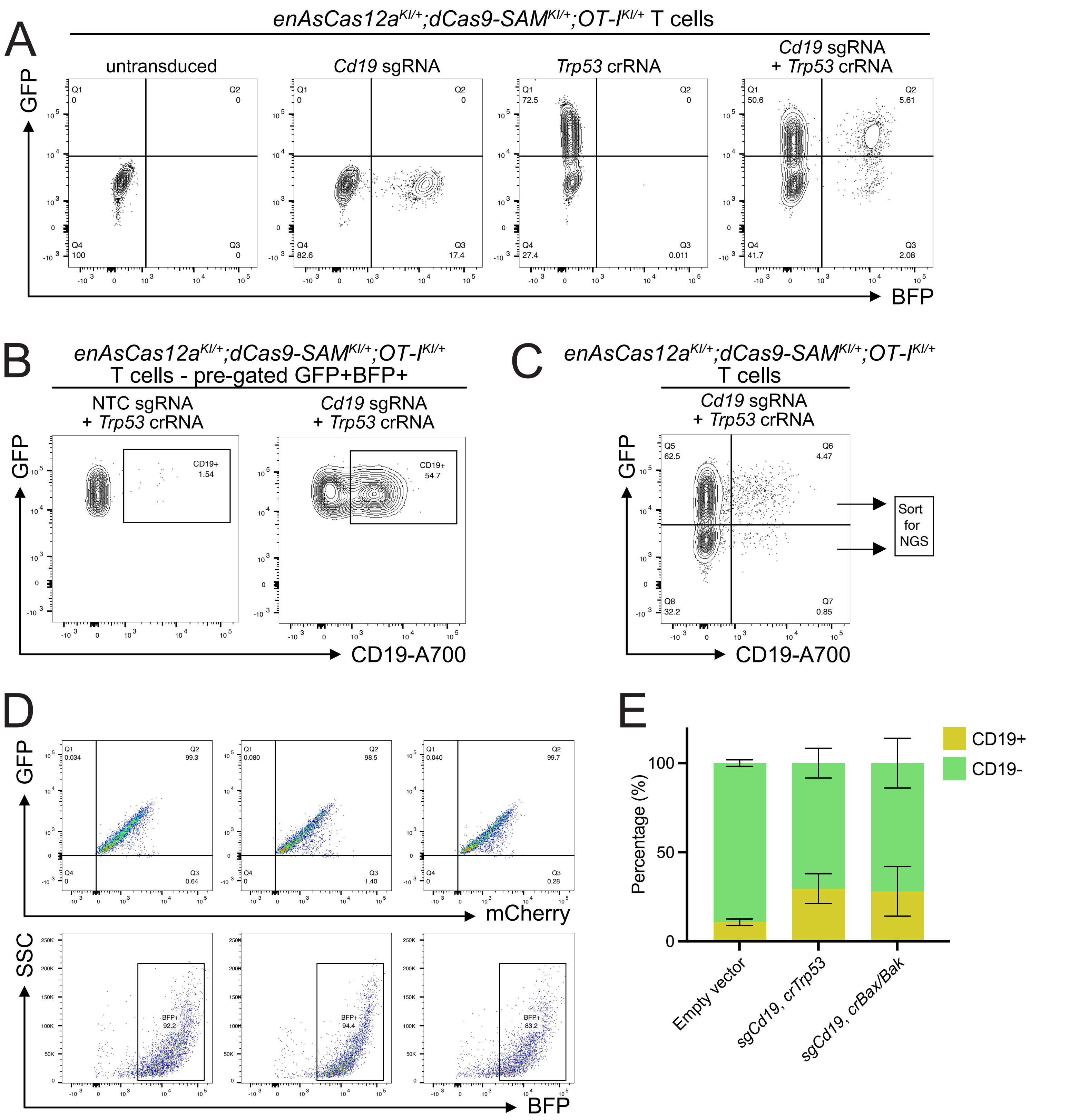
